# Regulation of spatial and temporal gene expression in an animal germline

**DOI:** 10.1101/348425

**Authors:** Asija Diag, Marcel Schilling, Filippos Klironomos, Salah Ayoub, Nikolaus Rajewsky

**Author notes:** These authors contributed equally to this work (Asija Diag, Marcel Schilling).

## Abstract

In animal germlines, regulation of cell proliferation and differentiation is particularly important but poorly understood. Here, using a cryo-cut approach, we mapped RNA expression along the *Caenorhabditis elegans* germline and, using mutants, dissected gene regulatory mechanisms that control spatio-temporal expression. We detected, at near single-cell resolution, > 10,000 mRNAs, > 300 miRNAs and numerous novel miRNAs. Most RNAs were organized in distinct spatial patterns. Germline-specific miRNAs and their targets were co-localized. Moreover, we observed differential 3’ UTR isoform usage for hundreds of mRNAs. In tumorous *gld-2 gld-1* mutants, gene expression was strongly perturbed. In particular, differential 3’ UTR usage was significantly impaired. We propose that PIE-1, a transcriptional repressor, functions to maintain spatial gene expression. Our data also suggest that *cpsf-4* and *fipp-1* control differential 3’ UTR usage for hundreds of genes. Finally, we constructed a “virtual gonad” enabling “virtual *in situ* hybridizations” and access to all data (https://shiny.mdc-berlin.de/spacegerm/).

## INTRODUCTION

Spatial and temporal restriction of gene expression has been considered for decades to be a crucial conserved mechanism for cellular and developmental programs such as specification of cell fates and compartmentalization. The function of mRNA localization is diverse (Buxbaum et al., 2015; Jansen, 2001; Martin and Ephrussi, 2009). On the one hand, localization of mRNA is thought to be more energy efficient as it serves as a template for multiple rounds of translation (Jansen, 2001; Martin and Ephrussi, 2009). On the other hand, local translation might protect other cells or compartments from proteins that are toxic for these cells or compartments (Martin and Ephrussi, 2009). Which mechanisms control mRNA localization? Recent studies suggested that alternative polyadenylation (APA) and hence 3’ Untranslated Regions (3’ UTRs) are an important post-transcriptional mechanism that regulates spatial restricted gene expression and cell fate transition (Brumbaugh et al., 2018; Mayr, 2017). Recent *in vivo* studies on the *C. elegans* germline revealed that 3’ UTR are the primary regulators of gene expression (Merritt et al., 2008). Furthermore, differential 3’ UTR usage can modulate the balance between proliferation and differentiation (Lackford et al., 2014; Mayr and Bartel, 2009; Sandberg et al., 2008; Shepard et al., 2011; Sood et al., 2006; 2009). Cells must decide, whether, when, where and how fast to proliferate in order to keep the balance between proliferation and differentiation as improper regulation can lead to developmental defects and cancer. However, our understanding of how mRNA localization regulates the balance between proliferation and differentiation remains limited.

The *C. elegans* germline is a powerful *in vivo* model for studying the balance between proliferation and differentiation. The basic factors, molecular architecture and processes are similar to that of other metazoans and major players have been remarkably conserved during evolution. The germline is divided into different compartments: In the distal portion of each arm and in close proximity to the germline niche (distal tip cell) proliferative germ cells are located, which form a syncytial tissue (Hirsh et al., 1976) (**Figure 1A**). In the distal arm, at a defined distance from the niche, germ cells exit the mitotic cell cycle and start differentiation by entering meiosis (**Figure 1A**). This switch from proliferation to differentiation is termed mitosis-to-meiosis transition. As part of the intrinsic oogenesis program, many early germ cells undergo apoptosis, around the bend region (Gartner et al., 2008). Only certain germ cells differentiate to become oocyte or sperm.

Previous studies already showed evidence that spatio-temporal restriction of RNA binding proteins (RBPs) in the *C. elegans* germline can regulate mRNA expression by binding to their 3’ UTR (Crittenden et al., 2006; Nousch and Eckmann, 2013). One important example is GLD-1, an RBP that binds multiple mRNAs including its own mRNA, thereby regulating the switch from proliferation to differentiation (Brenner and Schedl, 2016; Francis et al.; Jones et al., 1996; Jungkamp et al., 2011). Additionally, GLD-2, the cytoplasmic poly(A)-polymerase (cytoPAP) in the *C. elegans* germline, is accumulating around the pachytene stage in the germline, and promotes meiotic entry by polyadenylation of mRNAs that are required for differentiation (Millonigg et al., 2014; Nousch et al., 2014, 2017).

Besides RBPs and 3’ UTRs being key players in regulating mRNA stability, in translation and in localization, previous studies suggested that microRNAs (miRNAs) might control proliferation and differentiation in the *C. elegans* germline (Bukhari et al., 2012; Ding et al., 2008). MicroRNAs (miRNAs), belonging to the class of small non-coding RNAs are important and conserved post-transcriptional regulators of gene expression that bind mRNAs, primarily in their 3’ UTR (Bartel, 2018). Usually, miRNA binding leads to transcript destabilization and/or translational inhibition. Bukhari and colleagues showed that loss-of-function of *alg-1* and *alg-2*, two miRNA-specific Argonaute proteins in *C. elegans*, leads to a reduced mitotic region and less proliferative cells in the *C. elegans* germline, indicating an important role of miRNAs in controlling germ cell biogenesis in the germline (Bukhari et al., 2012). So far, due to technical limitations such as low RNA content of the *C. elegans* germline and lack of sequencing protocols for low input materials, it has not been possible to gain a system-wide spatio-temporal resolved characterization of miRNA expression during germ cell proliferation and differentiation, with exception of the oocyte-to-embryo transition (Stoeckius et al., 2014).

Here we established an optimized version of the tomo-seq approach (Junker et al., 2014) and improved sequencing protocols for the *C. elegans* germline. Thus, we were able to quantify mRNA and miRNA expression at near single cell resolution, as a function of position along the germline. We capture *in vivo* RNA expression during the entire development of germ cells through proliferation and differentiation. With our approach we were able to detect novel miRNAs with highly restricted expression. As we also analysed several mutants, our data offer specific insights into mechanisms which are functionally important during germ cell development. We compared the spatio-temporal resolved gene expression of wild type germline to the *gld-2 gld-1* double mutant germline, unravelling new potential key players such as PIE-1, a maternal protein that blocks transcription, in the transition from proliferation to differentiation. By careful bioinformatics analysis of our data we discovered also hundreds of novel 3’ UTRs which had escaped previous approaches probably because they were often specifically expressed. Furthermore, we discovered that widespread differential 3’ UTR usage takes place along the germline. Strikingly, this phenomenon, which is key for changing regulation of mRNAs across space and time, was perturbed in the *gld-2 gld-1* mutants. With the exception of *cpsf-4* and *fipp-1*, all other factors known to regulate alternative 3’ UTR usage were not perturbed in the mutant, strongly arguing that the dynamic expression of these two factors are key contributors to differential 3’UTR expression. To provide a user-friendly interface of our massive data and to analyse gene expression of different genes but compared to a “universal” germline reference coordinate system, we set out to create a 3D germline model. By collecting data from the literature, by mining our own microscopy data, and by mathematical modelling we were able to create “SPACEGERM”, a model that reflects germline gene expression at near single cell resolution. SPACEGERM can be interactively mined remotely via the internet and can be used to perform systematically “virtual *in situs*” for >10,000 mRNAs and hundreds of miRNAs. In summary, we created the first map of spatially and temporally resolved germline mRNA and miRNA expression and our analysis provides crucial insights into mechanisms and function of RNA during germline development.

## RESULTS

### mRNAs and miRNAs are localized in the germline

To investigate the spatial and temporal distribution of gene expression along the germline, we dissected and embedded gonads, which harbour the germline, of young adults in tissue freezing medium. This entire procedure takes only a few minutes, minimizing RNA degradation. We then cryo-sectioned the gonads into ~15 slices of 50 μm thickness and performed RNA-seq on each slice with slightly different experimental approaches for mRNAs and small RNAs (**Figure 1B**). Analysis of the mRNA data revealed that most reads matched known *C. elegans* transcripts. Furthermore, quantified transcripts were in line with a poly(A) selection profile as expected due to the barcoded oligo(dT) primer used for capture (**Figure S1A, S1B** and **S4A**). Comparing pairs of biological and technical replicates confirmed that our experimental approach is highly reproducible with a Pearson correlation coefficient of 0.96 (**Figure S1C** and **S4B**). Additionally, as a control, we performed our experimental approach with uncut gonads and compared it to a sliced gonad in order to investigate if the slicing had an influence on the measured gene expression. By averaging the measurement of gene expression across slices, we were even able to reconstruct *in silico* the gene expression profile of the uncut gonad (**Figure S1D** and **S4C**). Furthermore, we showed that our sequencing method is reliable, since it compared very well to other sequencing approaches such as Poly(A)+-seq and ribosomal RNA depleted total RNA-seq (**Figure S1F**). As our biological replicates were slightly shifted and compressed to each other due to different cutting start points, we aligned samples to a common coordinate system (see Methods) before integrating the data of all replicates for downstream analyses. The *C. elegans* hermaphrodite germline contains two gonad arms, the anterior and the posterior gonad arm. We cut both arms to investigate any difference in gene expression between the arms. However, as expected and in concordance with the literature, we did not detect any difference between both arms above background (**Figure S2A**). Observable differences in gene expression between anterior and posterior decrease with rising expression levels, arguing that these differences reflect noise. Hence, we treated the samples as biological replicates and could therefore increase the statistical power of our analysis. Investigating the expression of mRNAs and miRNAs, revealed that both RNA classes display distinct localization pattern across the germline (**Figure 1C, 1D**, **S2B**, **S2C** and **S2D**). The gene expression profiles were consistent with *in situ* hybridization images of the gonad that we performed (**Figure 1D, 1E,** **S2A**, **S2B** and **S2C**). Altogether, these results demonstrate that our sequencing approach is reproducible and reliable and that the data reveal spatio-temporal organization of mRNAs and miRNAs throughout the germline.

**Figure 1.**
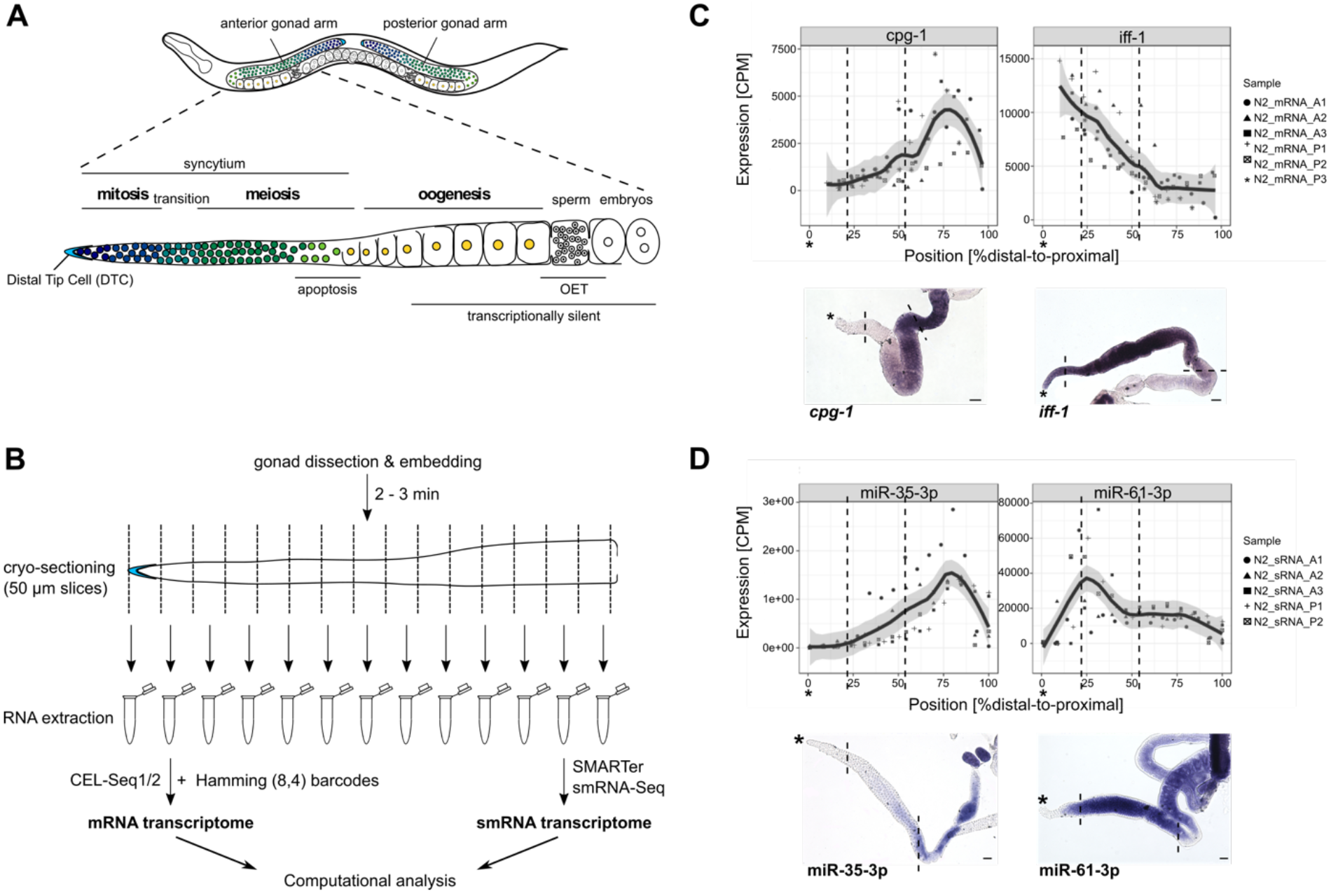
mRNAs and miRNAs are localized in the germline. (A) Schematic overview of the *Caenorhabditis elegans* gonad. OET: Oocyte-to-embryo transition. (B) Schematic overview of the experimental approach. (C) Spatial expression of *cpg-1* and *iff-1* from distal to proximal. n=6 independent experiments (N2_mRNA_A1-A3 and N2_mRNA_P1-P3) for wild type N2, LOESS ± standard error (SE). Corresponding *in situ* hybridization (ISH) images of *cpg-1* and *iff-1*. Asterisk: Distal tip cell (DTC). Scale bar: 20 μm. Dashed lines represent the different zones in the germline. (D) Spatial expression of mir-35-3p and miR-61-3p from distal to proximal. n=5 independent experiments (N2_sRNA_A1-A3 and N2_sRNA_P1-P2) for wild type N2, LOESS ± SE. Corresponding ISH images of mir-35-3p and miR-61-3p. Asterisk: DTC. Scale bar: 20 μm. Dashed lines represent the different zones in the germline. See also Figure S1, S2 and S4.

### A 3D germline model reflects RNA localization through germ cell proliferation and differentiation

As our data enables the systematic expression profiling of RNA along the germline in wild type and mutants, we thought that it would be useful to construct a model of the germline that can serve as a generalized framework on which expression data can be displayed and compared. We systematically collected published data about the size and composition of each zone in the germline (Brenner and Schedl, 2016; Fox et al., 2011; Hansen and Schedl, 2013; Hirsh et al., 1976; Hubbard, 2007; Maciejowski et al., 2006; Wolke et al., 2007) and quantified our own gonad images (**Table S1**). Using these data, we were able to compute an *in silico* 3D physical germline model (METHODS, **Figure 2A**). Within the model we assigned the number of germ cells to each zone and defined the size of each zone in the germline (**Figure 2A**). Despite the simplifications of the 3D model, we were able to reliably reveal RNA expression throughout the germline by integrating our sequencing data into the model (**Figure 2B, 2C** and **2D**). We used the 3D model as a guide to assign the different germline zones to our expression profiles. Thus, our 3D germline model integrates *in vivo* mRNA and miRNA expression throughout proliferation and differentiation of germ cells (**Figure 2B, 2C** and **2D**). Moreover, the model represents *in vivo* mRNA expression in perturbed systems such as the *gld-2 gld-1* double mutant (**Figure 2B**). In order to validate the assignment of the zones in our 3D model, we searched for apoptotic gene markers in the germline as most of the germ cells undergo apoptosis around the bend region. Indeed, we found apoptotic genes such as *ced-4*, having their highest expression precisely around the bend region that starts at a distance of approx. 350 μm from the distal tip cell (DTC) (**Figure 2C**) in accordance with the assignment of the bend region in our model. Finally, our 3D model also represents miRNA localization throughout the germline (**Figure 2D**). Overall, we believe that the virtual germline may serve as a reference to future studies.

**Figure 2.**
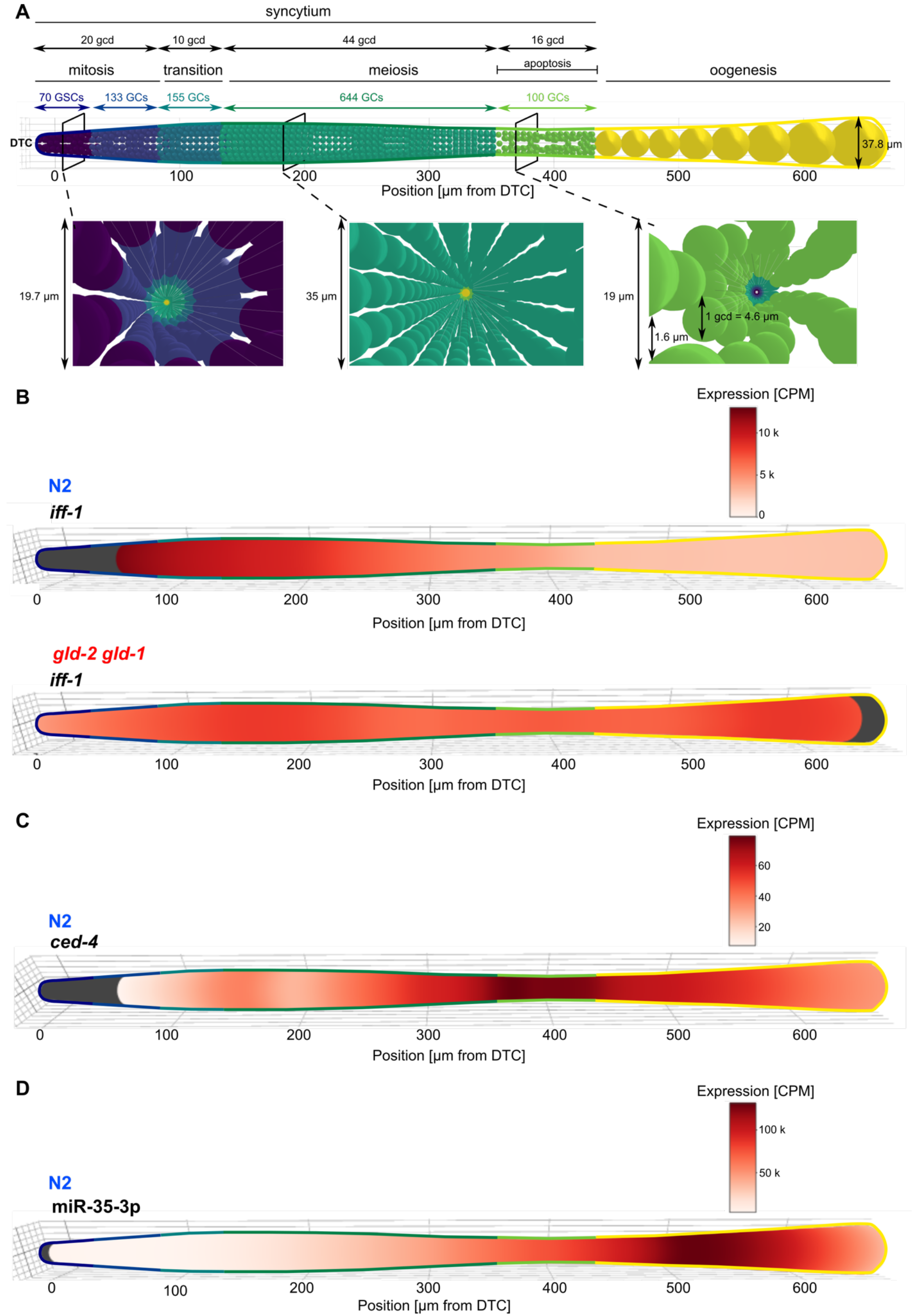
A 3D germline model reflects RNA localization throughout germ cell proliferation and differentiation. (A) 3D germline model with assigned sizes of each zone in germ cell diameter (gcd) and corresponding germ cell (GC) numbers. Three cross sections are shown at 70 μm, 200 μm and 380 μm form the distal tip cell (DTC). (B) 3D germline model representing *in vivo* expression of *iff-1* in N2 and *gld-2 gld-1* double mutant. Grey: No data. (C) 3D germline model representing *in vivo* expression of *ced-4* in N2. Grey: No data. (D) 3D germline model representing *in vivo* expression of miR-35-3p in N2. Grey: No data. See also Figure S7 and Table S1.

### Spatial gene expression is perturbed in *gld-2 gld-1* double mutants

To determine whether mRNAs display a global localization pattern throughout the germline we clustered the expression of germline specific genes (Wang et al., 2009) according to Pearson’s linear correlation (1 - Pearson’s r). Clustering the expression data revealed that mRNAs are organized in groups with distinct localization patterns (**Figure 3A**). We observed many different gene clusters along the germline. However, assigning these clusters to the zones in the germline (Brenner and Schedl, 2016; Hirsh et al., 1976) showed that most genes peaked in expression either in the mitotic or oogenesis region, whereas in the meiotic region genes required for proliferation/mitosis slowly decrease while genes required for differentiation/oogenesis slowly increase abundance.

To test whether mRNA localization is important for the transition between proliferation and differentiation we performed the cryo-based method for the *gld-2 gld-1* double mutant which possess only one third of meiotic entry (Brenner and Schedl, 2016; Kadyk and Kimble, 1998), ending up in a solely proliferating and tumorous germline. Hence, the *gld-2 gld-1* germline lacks oocytes and is sterile. Consistent with this fact, clustering the expression of germline specific genes for the *gld-2 gld-1* mutant revealed that mRNA localization is perturbed compared to the wild type (**Figure 3A**). In most cases, genes required for proliferation, *e.g.*, *iff-1*, were expressed continuously throughout the germline whereas genes required for embryogenesis were downregulated, *e.g.*, *perm-2* and *perm-4*. However, clustering of the same genes in the *gld-2 gld-1* double mutant, revealed that some genes still localize in the mutant (**Figure S3A**). In addition to the *gld-2 gld-1* double mutant, we further investigated the spatial gene expression of the *glp-1 (gf)* mutant germline which possesses a prolonged proliferative zone. The mutant is temperature sensitive resulting in an inducible tumorous phenotype. Hence dissection of gonads from these mutants was impeded. Therefore, we only induced the phenotype for a short time avoiding tumour development. As the induction of the mutation was very short, the spatial gene expression resembled more the wild type germline (**Figure S3B**).

Based on the finding that mRNAs are localized in the wild type germline and that this localization pattern is perturbed in the *gld-2 gld-1* mutant, we explored whether there are specific mRNAs localizing to a certain zone of the germline, *i.e.*, proliferation or differentiation. Investigating one specific cluster, distally peaking genes, in more detail, we observed that many genes encoding for a ribosomal subunit (*rpl* and *rps* genes) had their highest expression in the distal gonad arm and decreased in expression in the proximal arm (**Figure 3B**). Interestingly, this was not the case for the *gld-2 gld-1* double mutant (**Figure 3B**). The *rpl* and *rps* genes had a similar expression levels in the *gld-2 gld-1* double mutant as in the wild type but the expression did not decrease in the proximal arm but stayed constant along the germline, suggesting an important role of these genes in proliferation. This result was consistent with *in situ* hybridization images (**Figure 3C** and **3D**). In contrast to the *rpl* and *rps* genes, we observed some genes that had their highest expression in the proximal arm in the wild type while these genes were downregulated or completely absent in the *gld-2 gld-1* mutant (**Figure 3E**). We identified *pie-1, cey-2* and *nos-2* amongst these genes. The *pie-1* gene encodes for a maternal CCCH finger protein which is specific for oocytes and embryos (Merritt et al., 2008; Tenenhaus et al., 2001). Previous studies showed evidence that PIE-1 is a bifunctional protein that blocks the transcription of somatic transcripts during blastomere development, ensuring the germline fate and that it is required for the maintenance of class II mRNAs, mRNAs that are associated with P granules in the germline (Seydoux and Dunn, 1997; Seydoux and Fire, 1994; Seydoux et al., 1996; Tenenhaus et al., 2001). Additionally, it was shown that *pie-1* is a target of the cytoplasmic polymerase, GLD-2 (Kim et al., 2010). Consistent with these previous described findings, we observed in the *gld-2 gld-1* mutant, where GLD-2 is depleted, a strong downregulation of *pie-1* as well as *nos-2* and *cey-2*, two class II mRNAs. Together, these results suggest that *rpl* and *rps* genes are important for germ cell proliferation and that their transcription may be blocked by PIE-1 in the proximal arm while on the other hand *nos-2* and *cey-2* are important for differentiation which expression is maintained through PIE-1 expression.

**Figure 3.**
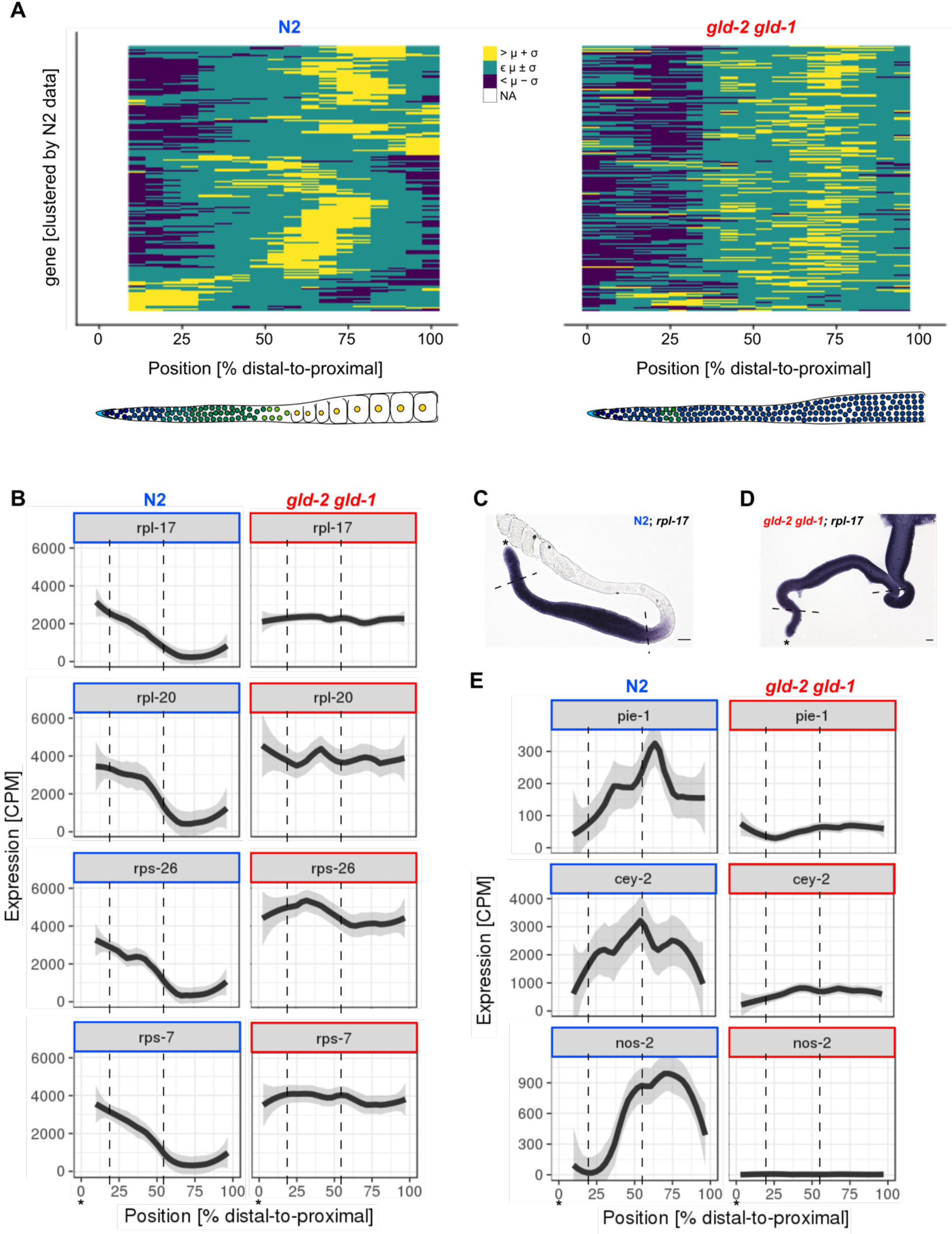
Spatial gene expression is perturbed in *gld-2 gld-1* double mutants. (A) Hierarchical clustering of germline specific genes by linear correlation (1 - Pearson’s r) for N2 and *gld-2 gld-1* double mutant. μ: Mean; σ. Standard deviation. NA: No data. (B) Spatial expression of two *rpl* genes (*rpl-17* and *rpl-20*) and two *rps* genes (*rps-26* and *rps-7*) in N2 and *gld-2 gld-1* double mutant from distal-to-proximal, respectively. n=6 independent experiments for N2 and n=4 independent experiments for *gld-2 gld-1* double mutant, LOESS ± standard error (SE). Dashed lines represent the different zones in the germline. Asterisk: Distal tip cell (DTC). (C) *In situ* hybridization (ISH) image of *rpl-17* in N2. Asterisk: DTC. Scale bar: 20 μm. Dashed lines represent different zones in the germline. (D) ISH of *rpl-17* in *gld-2 gld-1* double mutant. Asterisk: DTC. Scale bar: 20 μm. Dashed lines represent the different zones in the germline. Asterisk: DTC. (E) Spatial expression of *pie-1*, *cey-2* and *nos-2* in N2 and *gld-2 gld-1* double mutant from distal to proximal, respectively. n=6 independent experiments for N2 and n=4 independent experiments for *gld-2 gld-1* double mutant, LOESS ± SE. Dashed lines represent different zones in the germline. See also Figure S3.

### Germline-specific small RNA sequencing identifies novel miRNAs

In order to investigate the spatially restricted expression of miRNAs in the germline we used the SMARTer smRNA kit from Clontech^®^. The kit has a very low level of bias as adapter ligation is completely abolished (Dard-Dascot et al., 2018). Instead, the 3’ adapter is added after polyadenylation of the total RNA and by oligo-dT priming in that poly(A) tail and 5’ adapter is added through reverse transcriptase template-switching. A side effect of the SMARTer kit, as reported by Dard-Dascot and colleagues, is the high frequency of side products like adapter concatemers. However, standard preprocessing of small RNA sequencing raw reads requires efficient adapter trimming which removes such artefacts. While this approach has the potential to capture other small RNAs, we focused on miRNAs as this class of small RNAs was implicated in regulation of proliferation and differentiation during germline development (Bukhari et al., 2012). Of note, clustering the expression of all detected miRNAs in the germline revealed that miRNAs are organized spatially in the germline (**Figure 4A**). The spatial patterns of miRNAs were similar to those of mRNAs (**Figure 3A**).

Due to technical limitations, *i.e.*, sequencing of small RNAs with very low input material (≤ 1 ng total RNA), it is likely that miRNAs specifically expressed in the gonad or even limited to a specific region therein would have been missed by previous attempts to identify miRNAs as their signal would have been diluted out. As such highly specific miRNAs would be prime candidates for key regulators of spatial expression in the gonad, we screened our germline specific small RNA-seq data for potential novel miRNAs. Therefore, we ran miRDeep2 (Friedländer et al., 2012) on our data. Indeed, we were able to predict 83 novel precursor miRNAs (**Figure 4B** and **Table S2**). In order to quantify these novel miRNAs, we included the mature and precursors of the novel miRNA predictions in the miRBase21 reference and re-ran miRDeep2 for quantification of known and novel miRNAs in each slice separately. Novel miRNAs as well as known miRNAs were reproducibly quantifiable. Remarkably, most of the novel miRNAs were, when averaged over the germline, very lowly expressed (≤ 100 CPM), explaining why they were missed in previous studies. Unlike most known miRNAs, several novel miRNAs were primarily expressed in the distal part of the germline suggesting a specific role for these miRNAs in the proliferation (**Figure 4B**).

As low expression and distinct localization could be an indication of technical artefacts, we picked four (nov-1-3p, nov-63-3p, nov-72-3p and nov-82-5p) out of the 83 putative new precursor miRNAs with different expression levels for validation. All four candidates revealed a miRNA-like hairpin structure when folding their pre-miRNA sequence *in silico* with star and mature sequences extensively complementing each other (**Figure 4C**). Furthermore, the read coverage was miRNA-like with reads stacking up mostly on the mature sequence at aligned 5’ positions (**Figure S5A, S5B, S5C** and **S5D**). We were able to validate the expression of three novel miRNAs out of the four chosen ones either with TaqMan^®^ assay, an assay specific for small RNA detection, or with *in situ* hybridization experiments (**Figure 4D** and **4E**). Interestingly, nov-72-3p had a higher expression in the germline compared to let-7-5p and nov-63-3p revealed germline specificity as it was almost 8-fold enriched in the germline compared to the whole worm (**Figure 4D**). We could not validate nov-82-5p, maybe due to its low expression. Consistently, our analysis revealed that expression of miRNAs as measured by qPCR correlates well with the CPMs determined with our sequencing approach (**Figure S5E**) Interestingly, the loci of nov-63-3p, nov-72-3p and nov-82-5p were found covered by reads from DCR-1 PAR-CLIP data (Rybak-Wolf et al., 2014) and ALG-1 iPAR CLIP data (Grosswendt et al., 2014) previously published by our lab (**Table S2**) supporting the existence and functionality of these novel miRNAs.

Further, we investigated whether the novel miRNAs have distinct targets. For this propose, we used the miRNA:mRNA chimera data generated previously in our lab (Grosswendt et al., 2014). The chimera data was generated using L3 staged worms which lack a fully developed germline. Hence, we were not able to find almost any of our novel miRNA candidates in this data set. However, nov-72-3p revealed an interesting novel interaction pattern as it interacts not only with mRNAs but also with other already known miRNAs (**Figure S5**). The strongest interaction was discovered between nov-72-3p and miR-52-5p. The phenomenon of a miRNA-miRNA duplex was already described by Lai and colleagues in 2004 but lacked so far experimental evidence (Lai et al., 2004). Their hypotheses were that a miRNA-miRNA duplex could either protect the single stranded miRNA from degradation or that it tethers the miRNA away from its putative mRNA targets. However, we did not observe any significant positive correlation between nov-72-3p and corresponding miRNAs throughout the gonad (data not shown). As we were able to validate nov-72-3p also in whole worm samples, the interaction between nov-72-3p and other miRNAs may play an important role in other tissues of the worm rather than the germline. One main issue with nov-72-3p was that we were not able to trace back the origin of this novel miRNA as the 20 nt of mature mRNA mapped to the *dpy-2* locus whereas the first 17 nt also mapped to the locus of *rrn’s*. However, mapping to the *dpy-2* locus revealed that the seed region of nov-72-3p is strongly conserved amongst other species (**Figure S5C**).

We also found chimeric reads for nov-1-3p. Intriguingly, nov-1-3p and other miRNAs of the predicted novel ones harbour a stretch of A’s in the seed region, suggesting a potential novel class of miRNAs. We validated nov-1-3p with small RNA ISH as TaqMan^®^ probes cannot be designed against a stretch of A’s due to low complexity (**Figure S5G**). Overall, we detected a high fraction of novel miRNAs being localized throughout the germline.

**Figure 4.**
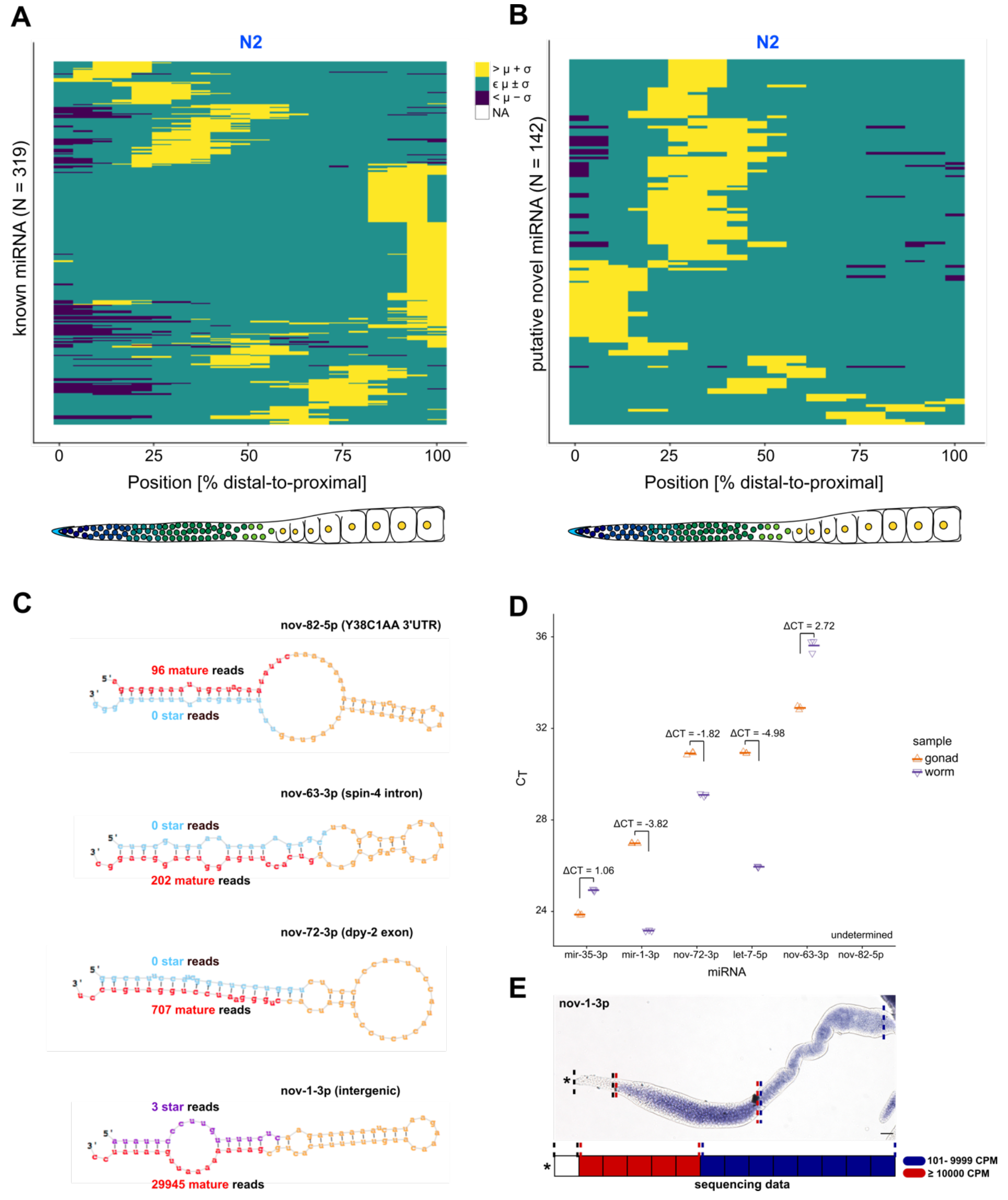
Germline-specific small RNA sequencing identifies novel miRNAs. (A) Hierarchical clustering of known miRNAs by linear correlation (1 - Pearson’s r) for N2. μ, mean; σ, SD. (B) Hierarchical clustering of novel miRNAs by linear correlation (1 - Pearson’s r) for N2. μ, mean; σ, SD. (C) Four examples of identified novel miRNAs of different *C. elegans* genomic origin. Reduced miRDeep2 plots show the precursor hairpin structure and the coverage of mature (red), star (blue, violet), and loop (yellow) sequences. (D) TaqMan^®^ assay validation (mean C_T_ values) of known miRNAs expressed in *C. elegans* (mir-35-3p, mir-1-3p and let-7-5p) and novel miRNA predictions (nov-63-3p, nov-72-3p and nov-82-5p). n=3 independent experiments for gonad and whole worm sample, respectively. (E) *In situ* hybridization (ISH) images of novel miRNA, nov-1-3p with corresponding spatial sequencing data. Asterisk: DTC. Scale bar: 20 μm. See also Figure S4, S5 and Table S2.

### A germline-specific miRNA family co-localizes with its putative targets

Based on the discovery that mRNAs and miRNAs revealed similar temporal and spatial expression patterns across germline, we asked whether expression of miRNAs and their corresponding targets is co-localized suggesting a putative miRNA:mRNA interaction. Instead of correlating each miRNA separately with putative targets, we correlated the family-wise expression summed among family members with the corresponding expression of the putative target. A miRNA family was defined by the 6mer seed found in the 2-7 nts of the miRNA members. Putative targets were identified by their 3’ UTR carrying at least one 7mer seed for the 2-8 nts of the miRNA or one 6mer seed for the 2-7 nts of the miRNA provided opposite the first miRNA nucleotide was an A (Bartel, 2009). We investigated the correlation for the miR-35 family members that are known to be germline-specific and that localize to the proximal gonad arm (**Figure 1D,** **2D** and **S2D**). Indeed, we showed that the miR-35-3p is enriched in the germline compared to whole worm (**Figure 4D**). The analysis revealed that all miR-35 family members in general correlate positively with their targets, *i.e.*, both classes showed co-localized expression throughout the germline indicating a germline-specific interaction (**Figure 5A**). In contrast, miR-1-3p, a non-germline specific miRNA that is expressed lowly in the germline compared to all miR-35 family members (**Figure 4D**), did not reveal any prominent co-localization pattern with its targets (**Figure 5A** and **5B**). As expected, miR-1-3p displayed anti-correlation with most of its targets implying an interaction outside of the germline. Overall our data suggest that germline-specific miRNAs co-localize with their targets which is a necessity for *in vivo* interaction.

**Figure 5.**
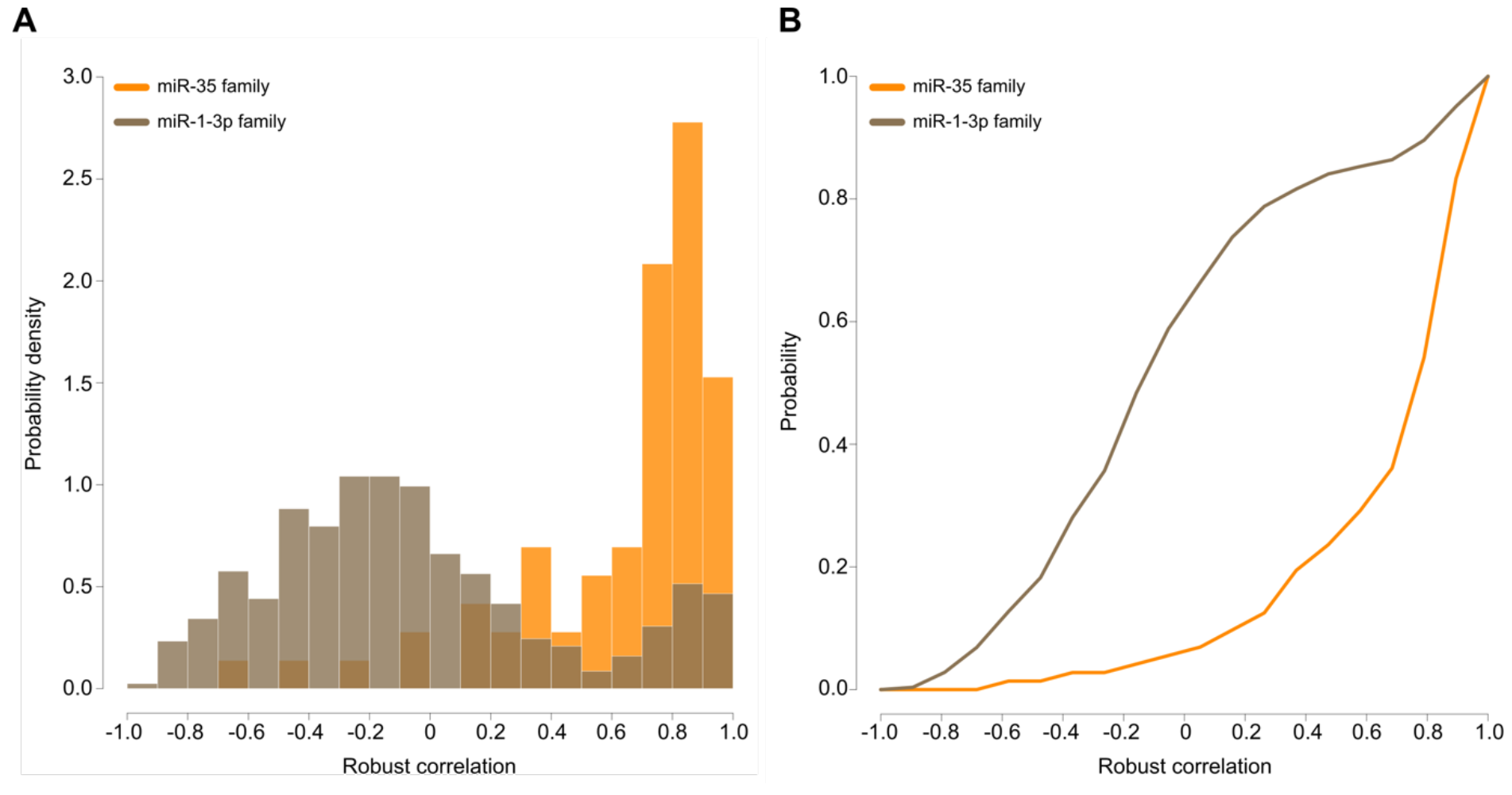
A germline-specific miRNA family co-localizes with its putative targets. (A) Histogram of robust correlation of miR-35 and miR-1 family members with their putative mRNA targets, respectively. (B) Density of robust correlation coefficients of miR-35 and miR-1 family members with their putative mRNA targets, respectively.

### Hundreds of novel 3’UTR isoforms detected in the germline, and hundreds of 3’UTRs are switched during development

Recent studies suggested that not promoters but 3’ UTRs are the main regulators of gene expression in the *C. elegans* germline (Merritt et al., 2008). Additionally, short 3’ UTRs are mainly expressed in proliferating cells whereas long 3’ UTRs are predominantly expressed in differentiating cells (Mayr and Bartel, 2009; Sandberg et al., 2008; Sood et al., 2006). Because the germline is divided in proliferating and differentiating cells we aimed to determine whether genes, expressing more than one isoform, change 3’ UTR length across germline. While most reads, as expected, map to genomic loci annotated as 3’ ends of protein coding genes, we observed several coverage peaks downstream of annotated genes, suggesting longer 3’ UTRs for these transcripts. Hence, we first extended the 3’ UTR annotation (METHODS and **Figure 5A**). We detected 499 intergenic peaks and assigned them to an upstream gene if the intergenic peak was less than 10 kb downstream (**Figure S6A** and **Table S3**). Of these intergenic peaks, we considered only the ones as novel 3’ UTRs that were less than 3 kb downstream of the assigned gene, leaving 419 candidates considered as novel 3’ UTRs (**Table S3**). Out of the 419 candidates we randomly picked ten and we were able to validate and confirm the identity of seven of them by Sanger-sequencing (**Figure S6B, S6C, S6D, S6E** and **S6F**). After annotation of 3’ UTRs and further downstream analysis, we quantified the change of the (relative) 3’ UTR usage along the germline for 910 genes. Interestingly, we observed that some of these genes used predominantly the distal polyadenylation signal (PAS) in the distal gonad arm (longer 3’ UTR) while the proximal PAS (shorter 3’ UTR) was used mainly in the proximal arm (**Figure 6B, 6C,** **S6G** and **S6H**). The switch occurred around the bend region of the gonad where almost 90 % of the cells undergo apoptosis (Hansen and Schedl, 2013). As miRNAs from the miR-35 family, a germline specific miRNA family, and other miRNAs have their highest expression around the bend region, *i.e.*, pachytene stage (**Figure 1D,** **S2D** and **4A**), it suggests that switching from distal to proximal PAS may be a mechanism to evade degradation of the transcript as longer 3’ UTRs usually harbour binding sites for miRNAs or other negative regulators.

### Differential 3’ UTR isoform usage is strongly perturbed in the *gld-2 gld-1* double mutant

To better understand the mechanism by which differential 3’ UTR usage occurs across the germline, we examined potential 3’ UTR switching candidates in the *gld-2 gld-1* mutant. Surprisingly, the mutant revealed much less candidates that switch isoform usage (**Figure S6I**). Moreover, the wild type switch of differential isoform usage was generally impaired in general in the *gld-2 gld-1* mutant (**Figure 6B, 6C, 6D, S6G, S6H** and **6I**). This switch did not occur in the *gld-2 gld-1* mutant but instead only the distal PAS was used throughout the germline. Based on the finding that the differential 3’ UTR usage takes place in wild type gonads and that it is perturbed in the *gld-2 gld-1* mutant, we explored whether factors involved in regulation of alternative polyadenylation (APA) are perturbed in the mutant, too. We investigated two factors in more detail, *fipp-1* and *cpsf-4. Fipp-1* is an ortholog of the human FIP1L1 and *cpsf-4* is an ortholog of the human CPSF4L. Both factors are components of the cleavage and polyadenylation (CPSF) complex that recognize the canonical PAS (AAUAA) and interact with the poly(A) polymerase and other factors, thereby inducing cleavage and polyadenylation (Kaufmann et al., 2004). Hence, both factors play a key role in the pre-mRNA 3’ end formation and APA. Interestingly, we observed that in the wild type both factors increased expression towards the proximal arm and peak around the bend region (**Figure 6E**). This is in line with the fact that the switch from distal to proximal PAS occurred around the bend region, too. Strikingly, the expression of both factors did not increase towards the proximal arm in the *gld-2 gld-1* mutant but stayed rather constant or increased slightly compared to the wild type (**Figure 6E**). This suggests that the level of these two factors may be important for the switch between distal and proximal PAS usage. Altogether our data showed that differential 3’ UTR usage is highly regulated in the germline and it indicates that the level of certain factors involved in APA may be important for the differential 3’ UTR usage.

**Figure 6.**
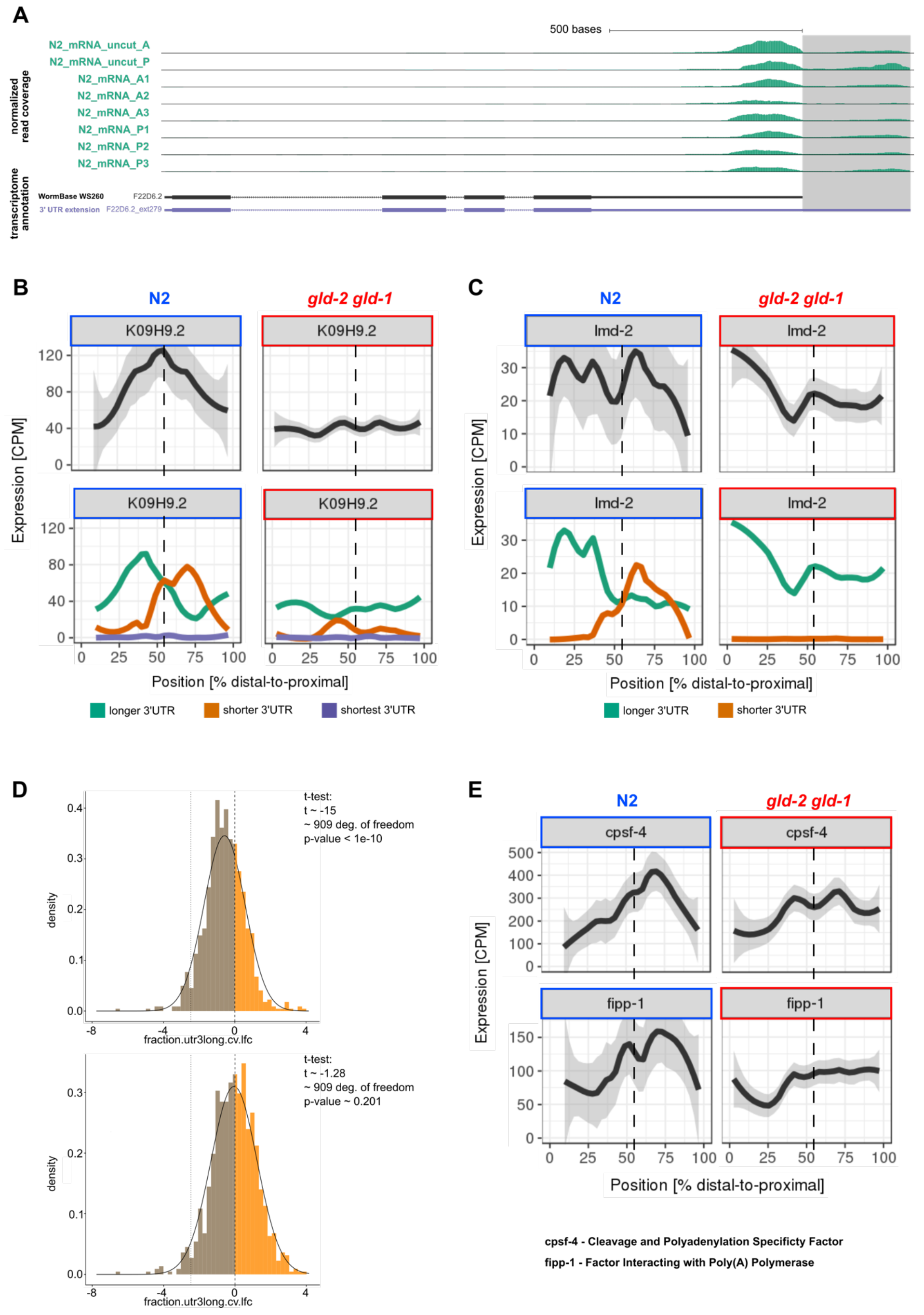
Differential 3’ UTR isoform usage in the germline is perturbed in the *gld-2 gld-1* double mutant. (A) Genome browser track example gene with downstream extension of the annotated 3’ UTR. (B) Spatial expression of K09H9.2 in wild type N2 and *gld-2 gld-1* double mutant from distal-to-proximal at gene and isoform level. n=6 independent experiments for N2 and n=4 for *gld-2 gld-1*, LOESS ± SE for gene level and LOESS only for isoform level. Longest 3’ UTR is marked in turquoise, shorter 3’ UTR in orange and the shortest 3’ UTR in purple. Dashed line, bend/loop region of the germline. (C) Spatial expression of *lmd-2* in wild type N2 and *gld-2 gld-1* double mutant from distal-to-proximal at gene and isoform level. n=6 independent experiments for N2 and n=4 for *gld-2 gld-1*, LOESS ± standard error (SE) for gene level and LOESS only for isoform level. Longest 3’ UTR is marked in turquoise and shorter 3’ UTR in orange. Dashed, bend/loop region of the germline. (D) Summary of log-CV (coefficient of variation) changes on a per-gene level between N2 and *gld-2 gld-1* double mutant. Lower panel shows the negative control with shuffled genotype assignments of the CVs. Left dashed line, the 5^th^ percentile of the shuffled control normal fit. (E) Spatial expression of *cpsf-4* and *fipp-1* in wild type N2 and *gld-2 gld-1* double mutant from distal-to-proximal. n=6 independent experiments for N2 and n=4 for *gld-2 gld-1*, LOESS ± SE. Dashed line marks the bend/loop region of the germline. See also Figure S6 and Table S3.

### SPACEGERM: a user-friendly interface for exploring spatial expression in the germline

We have shown how the spatially-resolved expression data generated in this study provide new insights into the mechanistic coordination of fundamental processes in biology. Clearly, these data have the potential to inform a multitude of additional studies focussing on various specific biological questions. To enable other researchers to conveniently utilize our data for their studies, and to provide a “universal” coordinate system, we developed SPACEGERM (<u>Spa</u>tial *<u>C.</u> <u>e</u>legans* <u>g</u>ermline <u>e</u>xpression of m<u>R</u>NA and <u>m</u>iRNA), an interactive data visualization tool for exploring the spatial expression data in the germline (**Figure S7**). The tool allows the user to investigate the spatial expression of every gene, isoform or miRNA detected in our data sets. The user can choose between wild type and mutant samples and have a closer look at the raw data points or the smooth fits (LOESS) across all replicates. SPACEGERM also allows to examine a set of genes by uploading an Excel file with gene name IDs, again for every genotype and gene type. Alternatively, one can investigate all genes detected with our sequencing approach up to 500 genes at once. Furthermore, the user can also download an Excel file with information about genes, their expression on average, their minimal and maximal expression value and location and cluster assignment. Finally, the reconstructed 3D germline can be explored concerning *in vivo* RNA expression throughout germ cell proliferation and differentiation (‘virtual *in situ* hybridization’ (vISH)).

## DISCUSSION

By rapidly dissecting, shock-freezing and cryo-cutting the *Caenorhabditis elegans* germline at 50 μm resolution and sequencing each slice separately, we create the first spatially resolved RNA expression map of wild type and mRNA expression map of mutant animal germlines. Additionally, we were able to reconstruct an *in silico* 3D germline model, that can be used to perform virtual *in situ* hybridizations and/or interrogate RNA localization of almost all transcripts during germ cell proliferation and differentiation (**Figure 2**).

### A mechanistic model of spatial gene expression regulation

We recovered the expression profile of *rpl* and *rps* genes, which encode for ribosomal subunits and are mainly localized to the distal gonad arm and slowly decrease in expression towards the proximal arm (**Figure 3**) (West et al., 2018). Interestingly, the same genes did not decrease in expression in the *gld-2 gld-1* double mutant but were constantly expressed throughout the germline (**Figure 3**). The *gld-2 gld-1* double mutant reveals only a third of the meiotic entry, *i.e.*, germ cells fail to differentiate and proliferate instead constantly throughout the germline (Brenner and Schedl, 2016; Kadyk and Kimble, 1998). However, genes involved in the deregulation of the proliferation and differentiation balance in the *gld-2 gld-1* mutant remain poorly discovered. In this study, we propose that PIE-1, a repressor of RNA polymerase II dependent gene expression that is important for germline cell fate determination (Seydoux and Dunn, 1997; Seydoux et al., 1996; Tenenhaus et al., 2001), is a potential key player that regulates the balance between proliferation and differentiation in the *C. elegans* germline (**Figure 7A**). The *pie-1* gene encodes a maternal CCCH finger protein which is specific for oocytes and embryos (Merritt et al., 2008; Tenenhaus et al., 2001). However, previous studies showed that the *pie-1* promoter allows expression in all germ cell types (D’Agostino et al., 2006; Merritt et al., 2008). Previous studies showed that *pie-1* is a target of GLD-2, the main cytoplasmic poly(A) polymerase (cytoPAP) in the germline (Kadyk and Kimble, 1998; Kim et al., 2010; Wang et al., 2009). Furthermore, Kim and colleagues showed that depletion of GLD-2 alone was sufficient to lower the abundance of most of its targets as these transcripts do not get polyadenylated and are therefore degraded (Kim et al., 2010). We showed that *pie-1* is mainly localized around the pachytene stage and that it has its highest expression during early oogenesis, exactly where the GLD-2 protein has its highest abundance (Millonigg et al., 2014). In accordance with this fact, the germline becomes transcriptionally silent from the late stage oogenesis (diagenesis) up to the forth cell-stage embryo (Evsikov et al., 2006; Stoeckius et al., 2014) suggesting that PIE-1 could play a key role in repressing the transcription as it does in the blastomere development. Indeed, many genes like the *rpl* and *rps* genes decrease in expression towards the proximal gonad arm supporting the hypothesis that PIE-1 is involved in transcriptional repression of these genes. In line with this hypothesis, when *pie-1* is downregulated in the *gld-2 gld-1* double mutant, *rpl* and *rps* genes are constantly expressed throughout the germline potentially inducing the observed phenotype of constant proliferation of germ cells. Additionally, previous expression studies of PIE-1 in HeLa cells reported that PIE-1 can inhibit transcription directly, suggesting a conserved mechanism (Batchelder et al., 1999). As PIE-1 is also detected in the cytoplasm, mainly in association with P granules (Mello et al., 1996; Tenenhaus et al., 2001), it was suggested that PIE-1 is required for the maintenance of *nos-2* and possibly other class II mRNAs, RNAs that are associated with P granules (Seydoux and Fire, 1994; Tenenhaus et al., 2001). Indeed, our data revealed that *nos-2* and *cey-2*, two examples of class II mRNAs, increase in expression as *pie-1* expression increases in the proximal arm in the wild type. The same mRNAs are down-regulated in the *gld-2 gld-1* mutant. This suggests that the downregulation of these class II mRNAs impede differentiation as the *gld-2 gld-1* mutant lacks differentiation which results in a sterile phenotype. Interestingly, RNA interference (RNAi) of *pie-1* revealed many phenotypes, including a sterile phenotype as in the *gld-2 gld-1* mutant (Melo and Ruvkun, 2012). However, meiotic entry and differentiation is only perturbed in *gld-2 gld-1* double mutants, *i.e*., meiotic entry appears normally in mutants lacking either *gld-1* or *gld-2* (single mutants) (Brenner and Schedl, 2016). Hence, we cannot exclude other regulators and the regulation via *gld-1* as most of the GLD-1 targets are downregulated in our data. Furthermore, the relationship between *gld-1* in its role in the distal meiotic entry decision and its role further down in the germline is complex as loss of *cye-1* and *cdk-2*, two important regulators of the mitotic cell cycle, causes the germline tumours still to differentiate (Fox et al., 2011).

### The choice of 3’ UTR is strongly regulated in the *C. elegans* germline

Merritt and colleagues already reported that 3’ UTRs and not promoters are the main drivers of gene expression in the germline (Merritt et al., 2008). Other studies also suggested that there might be a switch of 3’ UTR usage between proliferative and differentiating cells, more precisely, cells that proliferate use mainly the proximal alternative polyadenylation site (PAS, short 3’ UTR) while differentiating cells use predominantly the distal PAS (long 3’ UTR) (Mayr and Bartel, 2009; Sandberg et al., 2008; Sood et al., 2006). However, *in vivo* studies of differential 3’ UTR isoform usage remain still poorly investigated. Our sequencing approach does not offer the coverage and resolution to generally distinguish between different isoforms of one gene. This is because we sequence approximately 500 nt fragments while the mean 3’ UTR length of *C. elegans* transcripts is 211 nt (Jan et al., 2011; Mangone et al., 2010). Nonetheless, we succeeded in quantifying the change of the (relative) 3’ UTR usage along the germline for almost 1000 genes. It is important to note that due to our technical limitations, this number is almost certainly only a fraction of all 3’ UTR length switches.

Among these candidates, we observed genes that mainly used the proximal PAS in the distal gonad arm while the distal PAS was used in the proximal arm (**Figure 6** and **S6**). Our data revealed that *cpsf-4* (Cleavage and Polyadenylation Specificity Factor) and *fipp-1* (Factor Interacting with Poly(A) Polymerase), have their highest expression around the pachytene stage. This is exactly where the switch of differential 3’ UTR usage occurs (**Figure 6** and **S6**). Those two genes are known main regulators of APA in the *C. elegans* germline. They are probably the key spatial regulators, as other APA factors are lower abundant, less localized, and importantly, not perturbed in the mutants (see SPACEGERM). Furthermore, we discovered that these two factors are constantly expressed in the *gld-2 gld-1* double mutant leading to the failure of 3’ UTR length switching (**Figure 6** and **S6**). Additionally, the expression levels of the APA factors in the mutant were similar to those in the very distal gonad arm of the wild type. Our data suggest that the expression level of APA factors is crucial for the differential 3’ UTR usage and hence the switch between proliferation and differentiation during germline development. Lackford and colleagues already observed a similar phenomenon where alternative polyadenylation (APA) depends on the level of different factors involved in APA such as Fip1, an mRNA 3’ processing factor, and CPSF, a cleavage and polyadenylation specificity factor (Lackford et al., 2014). It is thought that generally the distal PAS is stronger than the proximal one, leading to the predominant usage of the distal PAS if the level of APA factors is low (Lackford et al., 2014). Thus, we propose that the spatial concentration of factors involved in APA are important for differential 3’ UTR usage along the germline and hence controlling proliferation versus differentiation (**Figure 7B**). Furthermore, the *gld-2 gld-1* mutant indicated that deregulation of the levels of factors involved in APA perturb the differential 3’ UTR usage and therefore may disturb the proliferation and differentiation balance. In general, the *gld-2 gld-1* mutant showed globally decreased 3’ UTR variability in the germline compared to the wild type. It remains still to be investigated what factors or pathways regulate the level of the factors involved in APA regulation and if mRNA expression level of APA factors mirrors the corresponding spatial protein expression. Furthermore, an approach is needed that increases the resolution to distinguish between different isoform of a gene in a spatial resolved manner. This will help to determine the total number of genes that follow our differential 3’ UTR usage hypothesis (**Figure 7B**).

**Figure 7.**
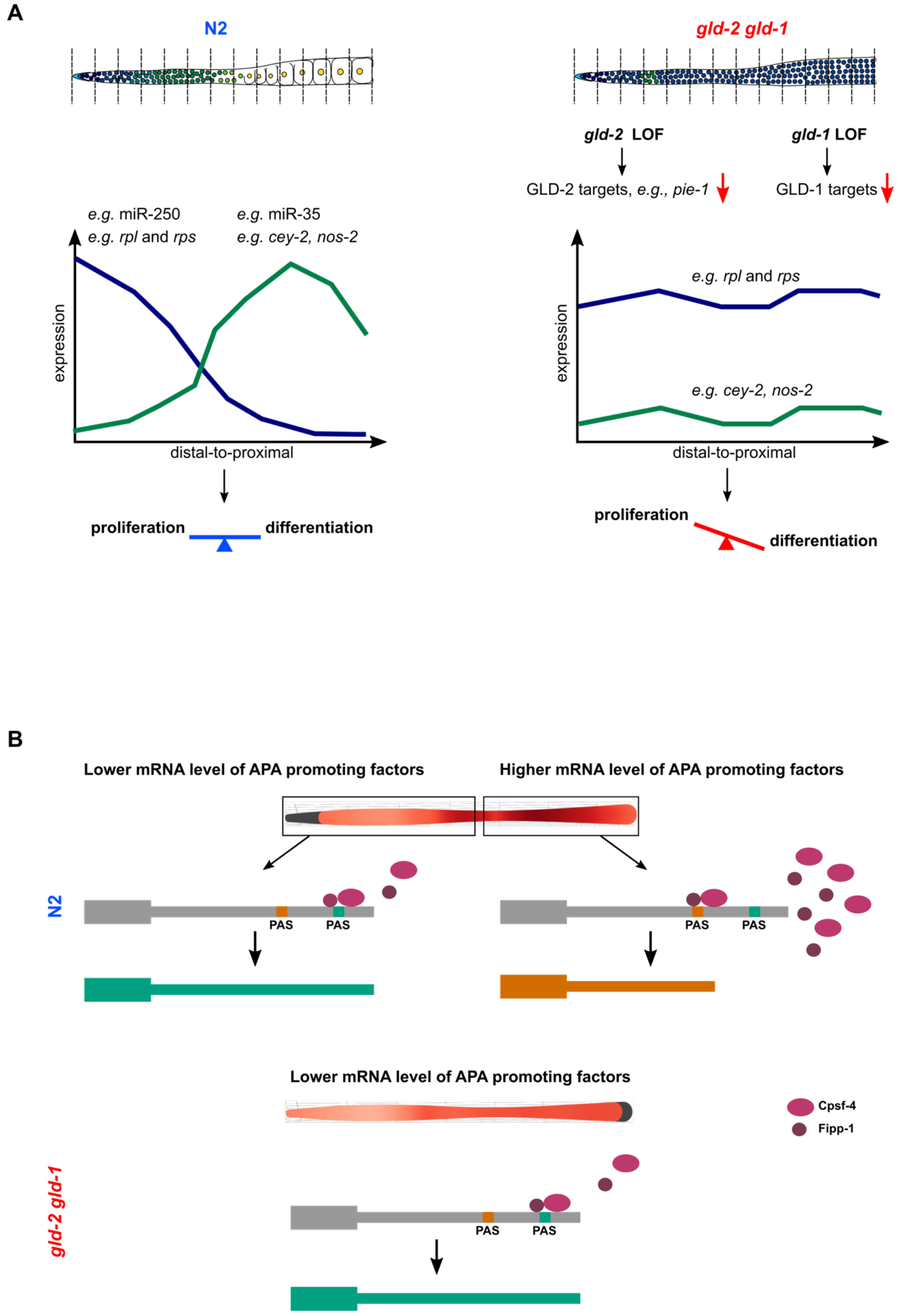
Model for spatially restricted gene expression and differential 3’ UTR isoform usage in the germline. (A) Schematic overview of mRNA and miRNA localization in wild type and mRNA localization in *gld-2 gld-1* double mutant germline, indicating putative regulators of spatial restricted gene expression. LOF, loss of function. (B) Model for differential 3’ UTR isoform usage across the germline. Depending on the concentration of *cpsf-4* (vISH is shown) and *fipp-1*, two factors involved in alternative polyadenylation (APA), some genes use the longer 3’ UTR isoform in the distal gonad arm while the shorter one is used in the proximal gonad arm. In *gld-2 gld-1* double mutants only the longer 3’ UTR isoform is used.

### miRNA expression is spatially organized and co-localizes with germline targets

Besides 3’ UTRs being important regulators of gene expression in the *C. elegans* germline, previous studies also suggested that miRNAs control proliferation and differentiation in *C. elegans* (Bukhari et al., 2012; Ding et al., 2008). We comprehensively analysed spatial expression of known miRNAs in the *C. elegans* germline. miRNAs were found in distinct localization patterns along the germline with the miR-35 family, the main miRNA family in the germline, being localized in the pachytene region (**Figure 1** and **4**), consistent with the previous study done by McEwan and colleagues (McEwen et al., 2016). Furthermore, we showed that all members of the miR-35 family and their targets were co-localized throughout the germline, a requirement for functional interaction *in vivo* (**Figure 5A**). One could speculate about a threshold function of the miR-35 family. In this case the miRNAs would keep the expression of their targets at a threshold level below which protein production is inhibited (Mukherji et al., 2011; Sood et al., 2006). In contrast, non-germline-specific miRNAs such as miR-1-3p did not show any predominant co-localization with their targets suggesting an interaction outside of the germline (**Figure 5A** and **5B**).

### Identification of 83 novel miRNAs with specific spatial localization

In addition, we discovered 83 novel precursor miRNAs and validated three of them (**Figure 4** and **S5**). We note that some of these miRNAs have a limited spatial expression domain but are well expressed within this domain. This is probably the reason why they escaped detection in previous studies. Interestingly, we identified an unusual novel miRNA, nov-72-3p, that (as we can show by chimera-analysis) binds other miRNAs, creating a miRNA-miRNA duplex (**Figure S5F**). This phenomenon was predicted by Lai and colleagues in 2004 computationally but so far lacked experimental evidence (Lai et al., 2004). Their hypotheses were that the miRNA-duplex could either stabilize the miRNA by protecting it from degradation or the miRNA could be tethered away from its targets and therefore stabilizing the mRNA targets (Lai et al., 2004). However, we were not able to define the genomic locus of nov-72-3p as the mature miRNA mapped antisense to the exon of the *dpy-2* locus but the first 17 nt of the mature miRNA also mapped to ribosomal RNA transcripts. Hence, the locus remains still undetermined impeding further analysis of nov-72-3p.

### Public availability of all data via interactive web application “SPACEGERM”

Finally, we developed an interactive data visualization tool, named SPACEGERM (<u>Spa</u>tial *<u>C</u>. <u>e</u>legans* <u>g</u>ermline <u>e</u>xpression of m<u>R</u>NA and <u>m</u>iRNA) for exploring the spatial expression in the germline in well defined, “universal” coordinates, both as raw data and projected (“virtual *in situ* hybridization”) on our 3D model (**Figure S7**).

Overall, we have presented a first map of germline RNA at unprecedented spatial resolution. This near single cell resolution was key for (1) discovering numerous of new miRNAs and hundreds of new 3’ UTRs (2) beginning to interpret the spatial patterns and (3) identifying, by comparison to mutant germlines, regulators and mechanisms that appear to play key roles in regulating germline biology. We believe that comparison to more mutants will dramatically improve our understanding of this beautiful system. Of course, much more measurements will need to be done as we currently only quantify RNA, and even for RNA we miss a lot of information – subcellular localization, methylation, polyadenylation states and many more. However, we hope that our 3D model and data help to set a common reference which can be expanded in the future.

## AUTHOR CONTRIBUTIONS

A.D. and N.R. conceived and designed the project. A.D. established and led the development of the project. A.D. performed the experiments and wrote (with N.R.) the manuscript, with input from other authors. M.S. analysed the mRNA data, designed the interactive data visualization tool and constructed the 3D germline model. F.K. analysed the small RNA data. S.A. helped with the establishment of the small RNA protocol. A.D. and N.R. led the interpretation of the data. N.R. supervised the project.

## ACKNOWLEDGMENTS

We thank J. P. Junker and B. Tursun for helpful discussions. We also thank the sequencing facility of the Sauer group at the BIMSB/MDC. We are grateful to M. Herzog for helping with the gonad dissection. We thank J. Hubbard for helpful discussions. We thank A. Filipchyk for identifying the chimeric interactions of nov-72-3p in the small RNA sequencing libraries. A.D. was a member of the Computational Systems Biology (CSB) graduate school which is funded by Deutsche Forschungsgemeinschaft (DFG). M.S. was funded by DFG and MDC. F.K. was funded by Deutsches Epigenom Programm (DEEP).

## METHODS

### Strains

All *C. elegans* strains were cultured by standard techniques (Brenner and Schedl, 2016). Worms were maintained at 16 °C on *E. coli* OP50-seeded nematode growth medium (NGM) plates. The following strains were used in this study: N2 Bristol wild type, *gld-2(q497) gld-1(q485)/hT2 [bli-4(e937) let-?(q782) qIs48]* (I;III) and *glp-1(ar202) III*.

### Embedding and cryo-sectioning

Gonads of wild type and mutants were dissected according to Francis and Nayak (Schedl lab) with minor modifications. The gonad, still attached to the worm body, was transferred to a specimen mold (Tissue-Tek^®^ cryomold^®^) filled with tissue freezing medium. This medium is very viscos, facilitating the stretching of the gonad and the separation from the worm body. Once the gonad was stretched, distal tip end and proximal end (at the end of oogenesis) were marked with AffiGel^®^ blue beads (Bio-Rad). Following, the specimen mold was rapidly frozen at −80 °C for 1 min and subsequently fixed in the cryotome to cut the gonad into slices of desired resolution. Each slice of the gonad was collected in an individual LoBind Eppendorf^®^ tube and immediately transferred to dry ice. RNA extraction of each slice was performed according Junker et al. (2014) with minor modifications. All experiments were performed in biological and technical triplicates for wild type and replicates for mutants for each gonad arm, *i.e.*, anterior and posterior gonad arm.

### mRNA library preparation

Reverse transcription and *in vitro* transcription were performed with the Ambion™ MessageAmp™ II kit according to CEL-seq method (Hashimshony et al., 2012) and tomo-seq method (Junker et al., 2014), except that all purification steps were performed using Agencourt^®^ AMPure^®^ XP beads according to CEL-seq2 (Hashimshony et al., 2016). Library preparation was performed using the Illumina TruSeq^®^ small RNA kit following the tomo-seq protocol (Junker et al., 2014). Unlike CEL-seq1/2 and tomo-seq, unanchored oligo(dT) barcodes used in this study were designed according to the Hamming [8,4] code allowing for barcode correction after sequencing (**Table S5**) (Bystrykh, 2012). For uncut samples and the first replicates of cut anterior and postrior gonad arm samples (N2_mRNA_A1 and N2_mRNA_P1), barcodes according to CEL-seq and tomo-seq (Hashimshony et al., 2012; Junker et al., 2014) were used. Libraries (with 30 % of PhiX spike-in DNA) were sequenced on the NextSeq 500 in a paired end mode.

### Small RNA library preparation

Small RNA libraries were only performed for the wild type N2 strain. Library preparation was performed for each slice separately using the SMARTer smRNA-Seq kit for Illumina from Clontech^®^ according to manufacturer’s instruction. Small RNA libraries of each slice were pooled and sequenced on HiSeq 2500 with a TruSeq^®^ 1 × 50 cycle kit as the Clontech^®^ kit is compatible with Illumina^®^ adapters and primers.

### Poly(A)+-selected library preparation

For the poly(A)+-selected library, several gonads were dissected and pooled. Library preparation was performed with the Illumina TruSeq^®^ stranded mRNA kit according to manufacturer’s instruction. Paired end sequencing was performed on the NextSeq 500.

### Ribosomal RNA depleted total RNA library preparation

For the ribosomal RNA depleted (ribodepleted) total RNA library several gonads were dissected and pooled. Ribosomal depletion was performed according to Adiconis *et al*. (2013). Library preparation was performed with Illumina TruSeq^®^ stranded total RNA kit. Paired end sequencing was performed on the NextSeq 500.

### Data pre-processing

Raw sequencing basecalls were demultiplexed and converted to FASTQ format using bcl2fastq v2.18.0.12 pooling reads across lanes (`--no-lane-splitting`). No adapter trimming was performed at this stage by not specifying adapter sequences in the sample sheet CSV file. To avoid masking of the short read 1 (barcode and UMI), the `--mask-short-adapter-reads=l0` option was used. 3’ reads (read 2) were annotated with their corresponding (corrected) barcode and UMI sequences (read 1) using custom scripts. Reads with identical barcode, UMI and sequence were collapsed and the unique reads were assigned to per-slice FASTQ files by barcode. Small RNA reads were subject to two rounds of 3’ end trimming by flexbar v. 2.5: The first round to remove 3’ adapters, the second to remove the poly(A)-tail added during the library preparation (using 10 A’s as ‘adapter sequence’). 3’ nucleotides with low basecall quality scores were trimmed using flexbars `--pre-trim-phred=30` option and the 3 nucleotides 5’ overhang introduced by the template-switching polymerase were trimmed using a custom awk script also discarding reads with a remaining length < 18 nts.

### Mapping of reads to the *C. elegans* genome

RNA-seq reads were mapped to the ce11/WBcel235 genome assembly using STAR_2.5.1b and an index with splice junction information from the Ensembl 82 transcriptome annotation. Alignments were sorted using sambamba v0.4.7. Coverage tracks were generated using bedtools v2.23.0 via the `genomecov` command specifying the `-split` and `-bg` options for splice-aware BedGraph output and splitting by strand using the `-strand` parameter. The total number of mapped reads per sample/splice was determined using the `flagstat` command of samtools 0.1.19-96b5f2294a and converted to the corresponding `-scale` parameter for `bedtools genomecov` for reads-per-million-mapped (RPM) normalization. Coverage BedGraph files were converted to BigWig format using bedGraphToBigWig v 4.

### 3’ extension of transcript annotation

The identification of downstream coverage peaks for 3’ extension of the WS260 transcriptome was performed using a custom R script: For each protein coding gene, the intergenic distance to the next downstream protein coding, ncRNA, lincRNA, pseudogene, rRNA or snoRNA gene (on the same strand) was calculated. Intergenic regions longer than 10 kb were truncated and the RPM-scaled genome coverage per sample of those regions was extracted from the BigWig files generated before. The per-sample coverage vectors were averaged per genomic position and binarized into uncovered regions (< 5 RPM mean coverage) and covered regions (>= 5 RPM mean coverage). Covered regions with a length of >= 50 nucleotides were considered as coverage peaks. Per downstream intergenic region, the downstream-most coverage peak was selected for the 3’ extension of the corresponding upstream gene. Only downstream extensions with a length up to 3 kb were considered for downstream analyses. For each gene with a downstream extension, all annotated transcript isoforms extending to the 3’ most genomic position of the corresponding gene were kept and got their 3’ UTRs extended by until the 3’ position of the respective downstream peak. Those 3’ extended transcripts were exported to a GTF file and merged with the WS260 transcriptome annotation using custom awk scripts.

### Transcriptome pre-processing

To enable the assignment of 3’ end RNA-seq reads to transcript isoforms, the 3’ extended WS260 transcriptome annotation was pre-processed using a series of custom R scripts: 3’ A’s were trimmed from all annotated transcripts as they would be indistinguishable from poly(A)-tails. The resulting transcripts were truncated to the 3’ most 500 nucleotides. Transcript isoforms with the same genomic coordinates and internal structure were collapsed and enumerated by decreasing corresponding (max.) 3’ UTR length.

### Isoform-specific transcript abundance estimation

RNA-seq reads were assigned to transcripts using kallisto 0.43.1: For 3’ reads, an index of the collapsed transcriptome annotation described above was used. For full-length coverage reads (poly(A)+ and ribodepleted total RNA-seq libraries), an index of the full 3’ extended transcriptome annotation was used. For all libraries, the `--bias` was passed to `kallisto quant`. For single end reads, additionally the `--single`, `--fragment-length=1` and `--sd=l` options were used. All libraries were sequenced with a first-strand-reverse stranded protocol. Thus, poly(A)+ and ribodepleted total RNA-seq samples were analyzed in `--rf-stranded` mode. The 3’ reads, while presented to kallisto as single-end reads, originally were sequenced as read 2, therefore resembling first-strand-forward single-end data. Thus, for these libraries the `--fr-stranded` mode of `kallisto quant` was used. Per-isoform read counts were exported to TSV files using the `--plaintext` option.

### Data processing

The raw read counts per transcript isoform and slice/sample were further processed using a custom R script: Though the whole annotated transcriptome was quantified to check for specificity of the experimental and computational approach, downstream analyses were limited to protein coding transcripts only. For gene-level analyses, isoform-level read counts were summed across all isoforms of a given gene. To compensate for differences in sequencing depth, raw read counts were normalized to counts-per-million (CPM). For full-length coverage protocols (poly(A)+ and ribodepleted total RNA-seq) an additional correction for the transcript length was performed, resulting in transcripts-per-million (TPM) estimates. Slice-data were arranged from distal to proximal by the known order of their barcodes and assigned to a relative position scale representing each slice by its center and accounting for differences in the number of slices per sample.

### Aligning cryo-cuts of different samples to a single coordinate system

As the start- and endpoint of gonad slicing was not precisely the same for all replicates, all slices of different replicates were aligned to a common coordinate system. This was achieved by comparing per-sample LOESS fits of abundance estimates across slices with *in situ* images of certain genes in the germline. Therefore, the gene profile of one replicate was fixed according to the corresponding *in situ* image and other replicates were aligned to the fixed replicate. This was done for approx. 30 gene profiles and the median of the shifting for those 30 profiles was calculated and used for all gene profiles.

### Integration of replicate data

The aligned discrete per-gene/isoform spatial expression profiles of individual replicates were used to fit a continuous consensus profile using local regression (LOESS) with a span of 0.4 through a custom R script. Slices with less than 10000 reads assigned to the transcriptome (‘dropout-slices’) were excluded from the fitting procedure. For visualization, 50 equidistant points along the distal-to-proximal axis were inferred from those fits. For downstream analyses, only 20 points were used to reflect the actual resolution of the data more conservatively. All data (incl. dropout-slices) are available through the interactive data exploration interface published alongside this study.

### Physical gonad model

To be able to assign the relative distal-to-proximal coordinates used for the spatially resolved gene expression profiles, a physical model of the *C. elegans* germline was built using a custom R script. The following assumptions were made for that model: i) Cells are approximately spherical. ii) Germ cells form a single layer tube within the distal part of the gonad arm. iii) The diameter of the gonad is minimal under the constraint of encompassing all germ cells. This enables a direct conversion between the number of cells in a germ cell layer and the diameter of that cell layer (given the size of a single germ cell) using basic geometry:

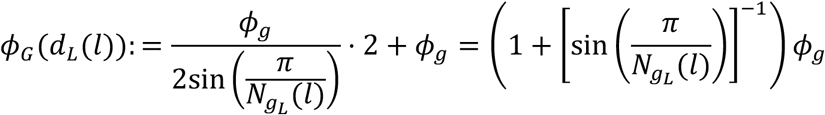

were *ϕ*_*g*_ is the diameter of a germ cell, *l* ∈ {1,10} ⊂ ℕ is the germ cell layer (one-based), 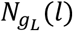 is the number of germ cells in layer *l*, *d*_*L*_(*l*) is the distance of the center of layer *l* to the distal tip cell (DTC) 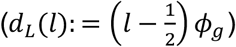 and *ϕ*_*G*_(*d*) is the diameter of the gonad arm at distance *d* from the DTC.

The (modelled constant) diameter of a single germ cell was set to 4.6 μm (Maciejowski et al., 2006). Based on our own measurements and results by Hirsh and colleagues (Hirsh et al., 1976) the total length of a stretched-out gonad arm was defined as 650 μm. At this distance to the distal tip cell (DTC) (*i.e.*, at the proximal end), the gonad must fit a fully mature oocyte, while at the distal-most end only a single germ cell needs to be fit in the gonad arm. To get a rough estimate of the size of a fully matured oocyte, the number of cells per embryo (558) (Wolke et al., 2007) was multiplied with the volume of a single germ cell. Given the equality in diameter of embryonic cells and germ cells and the equality in volume of the mature oocyte and the embryo, this gives a direct estimate for the size of the oocyte. To be able to model the gonad diameter in-between those extreme boundaries, we measured four gonad arms based on microscopic images (Table S1). Using these measurements at discrete points, a spline fit was used to model the radius of the gonad arm as a function of the distance to the DTC. Using this fit, the outline of the stretched-out gonad arm was modeled as a solid of revolution around the distal-to-proximal axis:

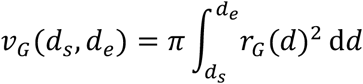

where *d*_*s*_ and *d*_*e*_ denote the distance to the DTC of the start and the end of the interval of interest, respectively, and *v*_*G*_(*d*_*s*_,*d*_*e*_) is the volume of the corresponding part of the gonad arm.

Based on the assumptions introduced above, the distal arm was filled with 1,002 germ cells in layers maximizing the number of cells per layer under the constraint given by the corresponding gonad diameter:

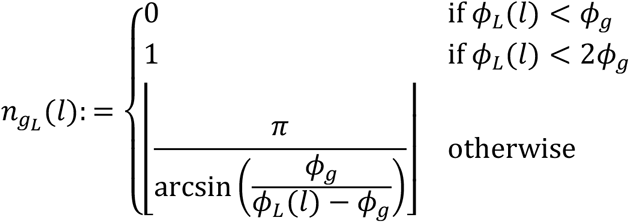

where 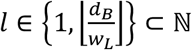 (*d*_*B*_ representing the distance to the DTC of the bend and *w*_*L*_: = *ϕ*_*g*_ the width of a germ cell layer) is the germ cell layer of interest (one-based), 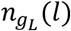 is the number of germ cells in that layer, and *ϕ*_*L*_(*l*) is the minimal diameter of the gonad in the interval containing germ cell layer *l*:

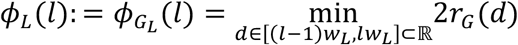

where *r*_*G*_(*d*) is the radius of the gonad at distance *d* from the DTC according to the spline model.

The total number of distal germ cells was derived from the total number of distal germ cell layers which was determined by comparing the cumulative number of cells up to each potential layer to the expected number of germ cells Ñ_*g*_: = 1000:

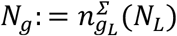

with

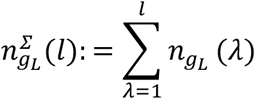

and

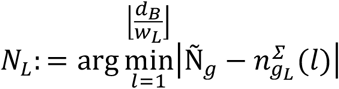

The mean distance in-between cells within the same layer resulting from this model was used as distance in-between germ-cell layers:

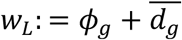

with

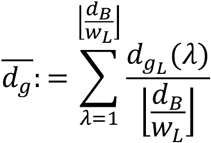

where

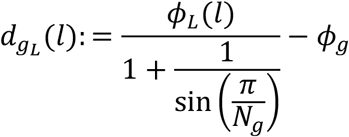

Germ layers were annotated functionally based on literature (Brenner and Schedl, 2016; Fox et al., 2011). The proximal gonad arm was filled with 8 oocytes, maximizing the diameter of each oocyte under the constraint of the corresponding gonad diameter. The proximal end of the distal germ cell layers and the distal end of the distal-most oocyte defined the boundaries of the loop region. Assuming steady-state with an apoptotic rate of 90% (Brenner and Schedl, 2016), the loop region was filled with 100 germ cells in layers (uniformly spread across the loop region).

### Data analysis of the small RNA transcriptome

The trimmed libraries were first mapped with bowtie2 (version 2.3.3.1) using the parameters `--very-fast-local --phred33 --local` to the *E. coli* genome (NC_000913.3, K-12, MG1655) in order to remove *E. coli* RNA contamination. The cleaned-up libraries were then mapped with STAR (version 2.5.3a) to the WBcel235/ce11 genome assembly using the Ensembl 87 annotation and the parameters

`-alignlntronMax 140000 -alignSJDBoverhangMin 17`
`--alignSplicedMateMapLmin 30 --outFilterMultimapNmax 5`
`--outFilterMismatchNmax 2 --outFilterMatchNmin 17`
`--outFilterScoreMinOverLread 0 --outFilterMatchNminOverLread 0.`

Sense and antisense read counting on features was done using HTSeq (version 0.9.1) with the parameters `-a 0 -m intersection-nonempty --nonunique=all --secondary-alignments=score` combined with `-s yes` for sense and with `-s reverse` for antisense counts. Known and novel miRNAs were identified separately using the cleaned-up libraries and the miRDeep2 algorithm (version 2.0.0.7) with the miRBase21 reference. First, miRDeep2 was ran on the pooled libraries. Then, the novel miRNA predictions found were added to the miRBase21 reference. Consequently, the combined reference of known and novel miRNAs was used for a second run of miRDeep2 on each library separately and on the pooled library as well. This way we unified the expression estimates of known and novel miRNAs under a common measure of counts per million of mapped reads (CPM).

The miRNA-target correlation analysis used robust linear regression based on the MM-estimator in order to reduce the effect of outliers (Koller and Stahel, 2011). All miRNAs were divided into families based on their 2 - 7nt 6mer seeds (reverse-complemented). Putative target genes were identified by counting miRNA 7mer seeds on all of their unique and longest 3’ UTR isoforms. The 3’ UTR isoform with the maximum number of 7mer seeds was taken as representative for that miRNA-target gene interaction. The miRNA 7mer seeds were chosen to be either the reverse-complement of the miRNA 2 - 8nts or the reverse-complement of the miRNA 2 - 7nts immediately followed by an A (Bartel, 2009). The control list of targets was generated by mutating the 3^rd^ and 4^th^ nucleotides of these 7mer seeds. Robust linear regression was done by summing the LOESS smoothed CPMs among the miRNA family members on each LOESS point and using this summarized family-wise expression with the corresponding target smoothed expression. In order for a correlation to be considered we demanded that both the family-wise miRNA expression and the target expression were commonly non-zero in at least 25% of the LOESS points.

### Probe preparation for mRNA in situ hybridization (ISH)

Digoxigenin (DIG)-labeled anti-sense RNA probes were prepared by *in vitro* transcription using a PCR generated DNA template. PCR primers were designed using Primer3 to amplify a 300-500 nt fragment from the cDNA prepared from whole worm samples. The T7 promoter sequence was added to the reverse primer to produce later an antisense probe by *in vitro* transcription. Primer sequences are provided in **Table S4**. PCR fragments were cleaned-up using Agencourt^®^ AMPure^®^ XP beads and *in vitro* transcription was performed with 0.5 - 1 μg DNA template using the T7 RNA polymerase and a DIG-RNA labeling mix (Invitrogen). Remaining DNA template was digested with DNase I and the RNA probe was precipitated with sodium acetate and ethanol for at least 30 min at −80 °C. After centrifugation the RNA pellet was washed with 75 *%* ethanol and probe integrity was checked on an agarose gel. The concentration of each RNA probe was adjusted to 50 ng/μl using 10 mM Tris-HCl/formamide solution (1:1).

### mRNA ISH

Worms were washed several times in sperm salt buffer only (100 mM PIPES, pH 7.0; 90 mM NaCl; 50 mM KCl; 40 mM CaCl_2_; 20 mM KH_2_PO_4_) and in the final step in sperm salt buffer containing levamisole. Up to 15 worms were transferred to a poly-L-lysine coated slide containing 8 μl of sperm salt and gonads were dissected according Francis and Nayak (Schedl lab) with minor modifications. After dissection 8 μl of 4 % paraformaldehyde (PFA) were added to the dissected gonads, a cover slip was put on top and the slide was incubated for 2 min. Following, the slide was incubated on dry ice for at least 20 min and the coverslip was flipped away using a razor blade under the coverslip (freeze and crack method). Slides were immediately immersed in ice-cold 100 % ethanol for 2 min, rehydrated in an ethanol series (90 %, 70 %, 50 %, 20%), following washing with PBS containing 0.2 % Tween for 30 min. Permeabilization of gonads was achieved with proteinase K treatment (1 μg/ml) for 5 min. Slides were washed in PBS containing 0.1 % Tween (PBS-T), fixed for 20 min in 4 % PFA, washed again with PBS-T, incubated in TEA buffer (aqua dest. containing 1.3 % triethanolamine; always prepared fresh), following final washing steps in PBS-T. Slides were prehybridized in prehybridization buffer (10 mM HEPES, pH 7.5; 600 mM NaCl; 50 m DTT; 1 mM EDTA; 1 × Denhardt’s solution; 100 μg/ml tRNA; 50 % formamide) for 1 h at 50 °C. Slides were hybridized over night at 50 °C in hybridization buffer (prehybridization buffer containing 10 % dextran sulphate) containing 0.5 - 1 μg/ml denaturated DIG-labeled antisense RNA probe (denaturation at 95 °C for 10 min). Slides were washed at 50 °C for 10 min with following solutions: posthybridization buffer (posthyb, 1 × 50 % formamide in 5 × SSC); 75 % posthyb buffer + 25 % 2 × SSC, 0.1 % Triton X; 50 % posthyb buffer + 50 % 2 × SSC, 0.1 % Triton X; 25 % posthyb buffer + 75 % 2 × SSC, 0.1 % Triton X; 2 × SSC, 0.1 % Triton X; 0.22 × SSC, 0.1 % Triton X. Following, slides were washed in maleic acid buffer (11.6 g/l maleic acid; 9.76 g/l NaCl; 0.1 % Triton X; pH 7.5) and afterwards incubated in 1 % blocking solution (Roche) diluted in maleic acid buffer for 1 h. Slides were incubated in Anti-DIG-AP (Roche, 1:2500) over night at 4 °C. After several washes with maleic acid buffer and TMN buffer (0.1 M Tris-HCl, pH 9.5; 0.1 M NaCl; 50 mM MgCl_2_; 1 % Tween 20; always prepared fresh), the signal was developed using NBT/BCIP (diluted in TMN buffer) solution. Time of development depended on the expression of the corresponding RNA and took up to 24 h for very lowly expressed RNAs. The background was removed with dehydration and rehydration in an ethanol series (samples were fixed before in 4 % PFA for 20 min again). For mounting, some μl of prolong gold (Invitrogen) were dropped on a coverslip and then inverted onto the slide. The edges were sealed with a nail polish.

### small RNA ISH

Gonad preparation and the prehybridization procedure was the same as for mRNA ISH. The TEA buffer contained additionally 0.06 N HCl and 0.27 % acetic anhydride. For small RNA ISH, DIG-labeled LNA (Locked Nucleic Acid) probes (former: Exiqon, now: Qiagen) were used (**Table S4**) and the prehybridization and hybridization temperature was set according to manufacturer’s instruction (20 - 25 °C below the melting temperature of the LNA probe). LNA probes were denaturated at 95 °C for 1 - 5 min prior hybridization. Prehybridization (without probe) was done for 1 h and hybridization (with 10 - 25 nM of LNA probe) over night. Slides were washed several times with 2 × SSC buffer and with 0.2 × SSC buffer. Following, the slides were washed with PBS-T and incubated for 1 h in blocking solution (PBS-T containing 5 % normal goat serum). Slides were incubated in Anti-DIG-AP (Roche, 1:2000) over night at 4 °C. After several washes with PBS-T and TMN buffer signal developing, background removal and mounting was performed according to mRNA ISH.

### TaqMan^®^ assays

The TaqMan^®^ assay was used to validate some of the novel miRNA predictions. TaqMan^®^ probes were designed with the Custom TaqMan^®^ Small RNA Assay Design Tool (ThermoFisher). TaqMan^®^ assays were performed according manufacturer’s instruction for gonad and whole worm samples. TaqMan^®^ target sequences are provided in **Table S4**.

### Nested PCR

Nested PCR was performed according to the Cold Spring Harbor Protocols (Sambrook and Russell, 2006). 0.5 - 1 μg of whole worm and gonad RNA were used as input RNA for the cDNA synthesis using TAP-VN as a primer. For the first nested PCR, 4 μl of 1:5 diluted cDNA was used. The first PCR was performed with a gene-specific forward primer and AP as a reverse primer. PCR products were purified using Agencourt^®^ AMPure^®^ XP beads and 10 - 20 ng of purified PCR were used for the second nested PCR. The second PCR was performed with a second gene-specific primer and MAP as a reverse primer. Annealing temperature was calculated using the NEB Tm Calculator (BioLabs). The PCR products from the second PCR were separated by agarose gel, purified and Sanger-sequenced to confirm the identity of the bands. Nested PCR was used for 3’ UTR extension validation. Alternatively, conventional PCR by designing the forward primer in the second last exon (to distinguish from genomic DNA) and the reverse primer in the 3’ UTR extension was used for validation (using whole worm RNA only). Primer sequences are provided in **Table S4**.

## SUPPLEMENTARY FIGURES

**Figure S1.**
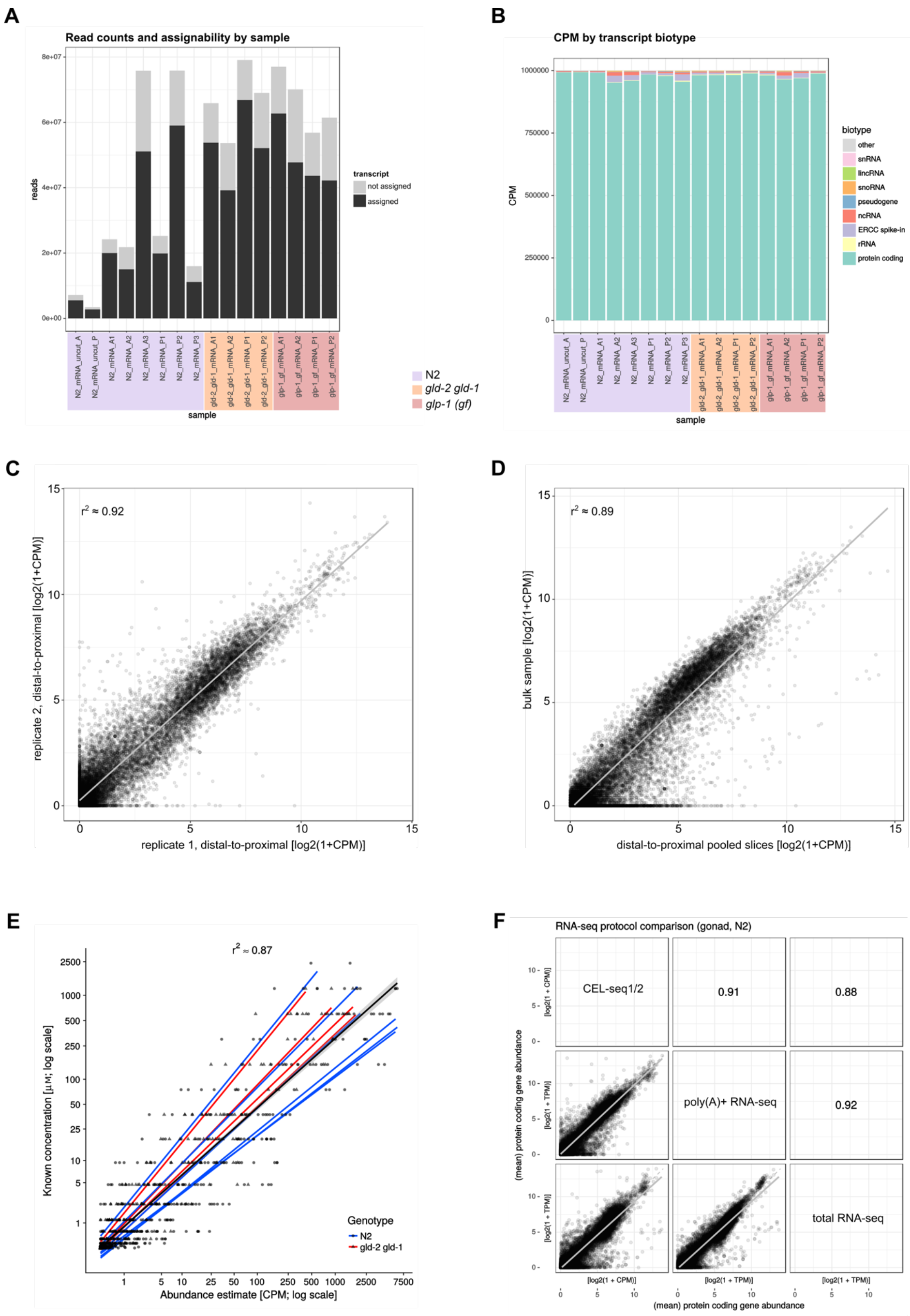
Experimental approach for spatial gene expression is reproducible and reliable. Related to Figure 1. (A) Read counts and assignability of reads for each biological and technical replicate of N2, *gld-2 gld-1* double mutant and *glp-1 (gf)* mutant. (B) Transcript biotype distribution over the fraction of mapped reads for each biological and technical replicate of N2, *gld-2 gld-1* double mutant and *glp-1 (gf)* mutant. (C) Linear correlation (Pearson’s r) across all transcripts, summed and averaged over all sections for two biological replicates. (D) Linear correlation (Pearson’s r) across all transcripts of uncut (bulk) sample and sliced samples (summed and averaged over all sections for all biological replicates). (E) Linear correlation (Pearson’s r) of known ERCC spike-in concentration and estimated spike-in abundance for N2 (red line) and *gld-2 gld-1* double mutant (blue line). (F) Linear correlation (Pearson’s r) across all genes for different sequencing approaches, *i.e.*, CEL-seq1/2, poly(A)+ RNA-seq and total RNA-seq.

**Figure S2.**
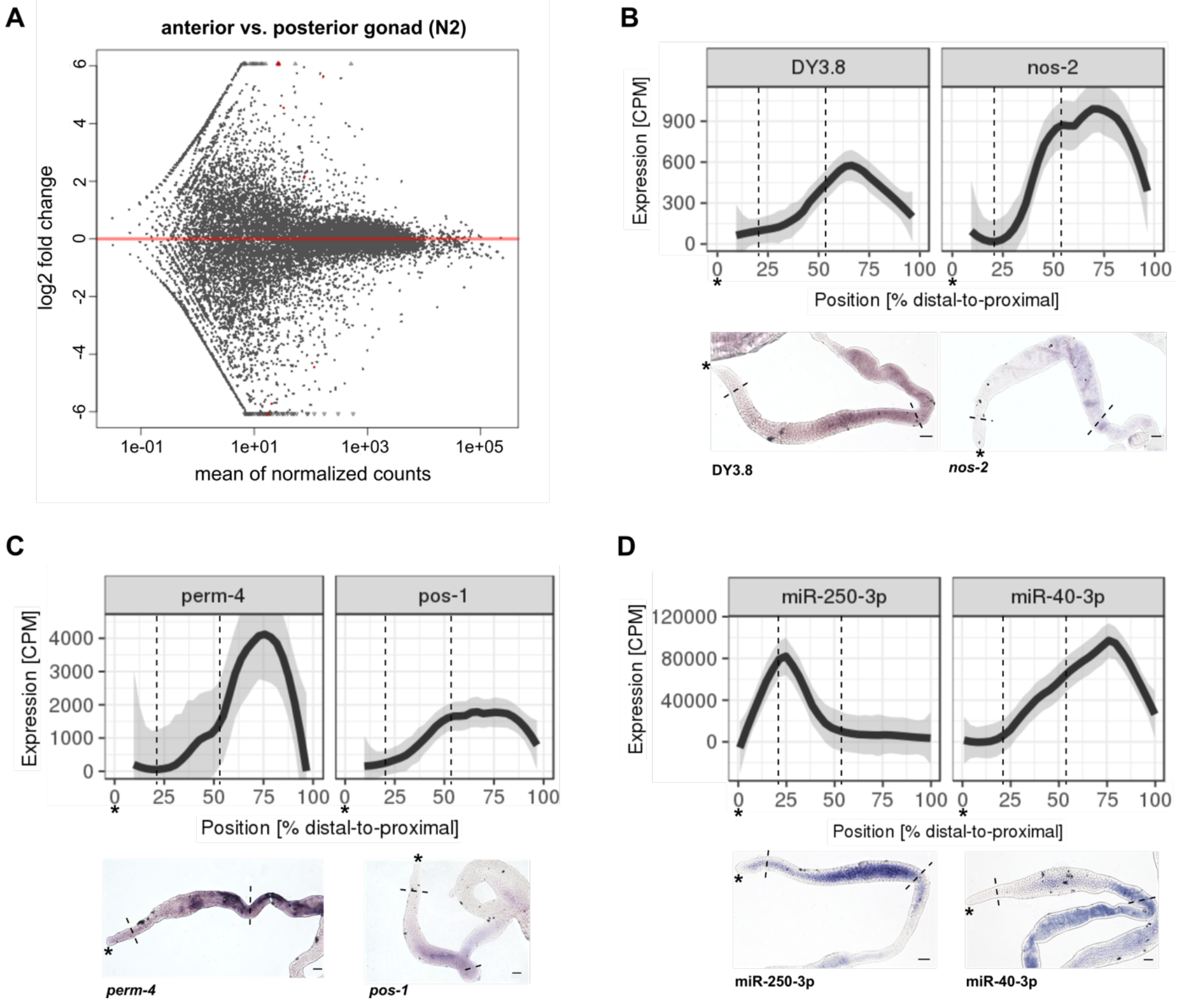
mRNAs and miRNAs are localized in the germline. Related to Figure 1. (A) Comparison of all N2 samples on the gene level by gonad arm (anterior or posterior) using DESeq2. (B) Spatial expression of DY3.8 and *nos-2* from distal to proximal. n=6 independent experiments, LOESS ± standard error (SE). Corresponding *in situ* hybridization (ISH) images of DY3.8 and *nos-2*. Asterisk: Distal tip cell (DTC). Scale bar: 20 μm. Dashed lines represent the different zones in the germline. (C) Spatial expression of *perm-4* and *pos-1* from distal to proximal. n=6 independent experiments, LOESS ± SE. Corresponding ISH images of *perm-4* and *pos-1*. Asterisk: DTC. Scale bar: 20 μm. Dashed lines represent the different zones in the germline. (D) Spatial expression of miR-250-3p and miR-40-3p from distal-to-proximal. n=6 independent experiments, LOESS ± SE. Corresponding ISH images of miR-250-3p and miR-40-3p. Asterisk: DTC. Scale bar: 20 μm. Dashed lines represent the different zones in the germline.

**Figure S3.**
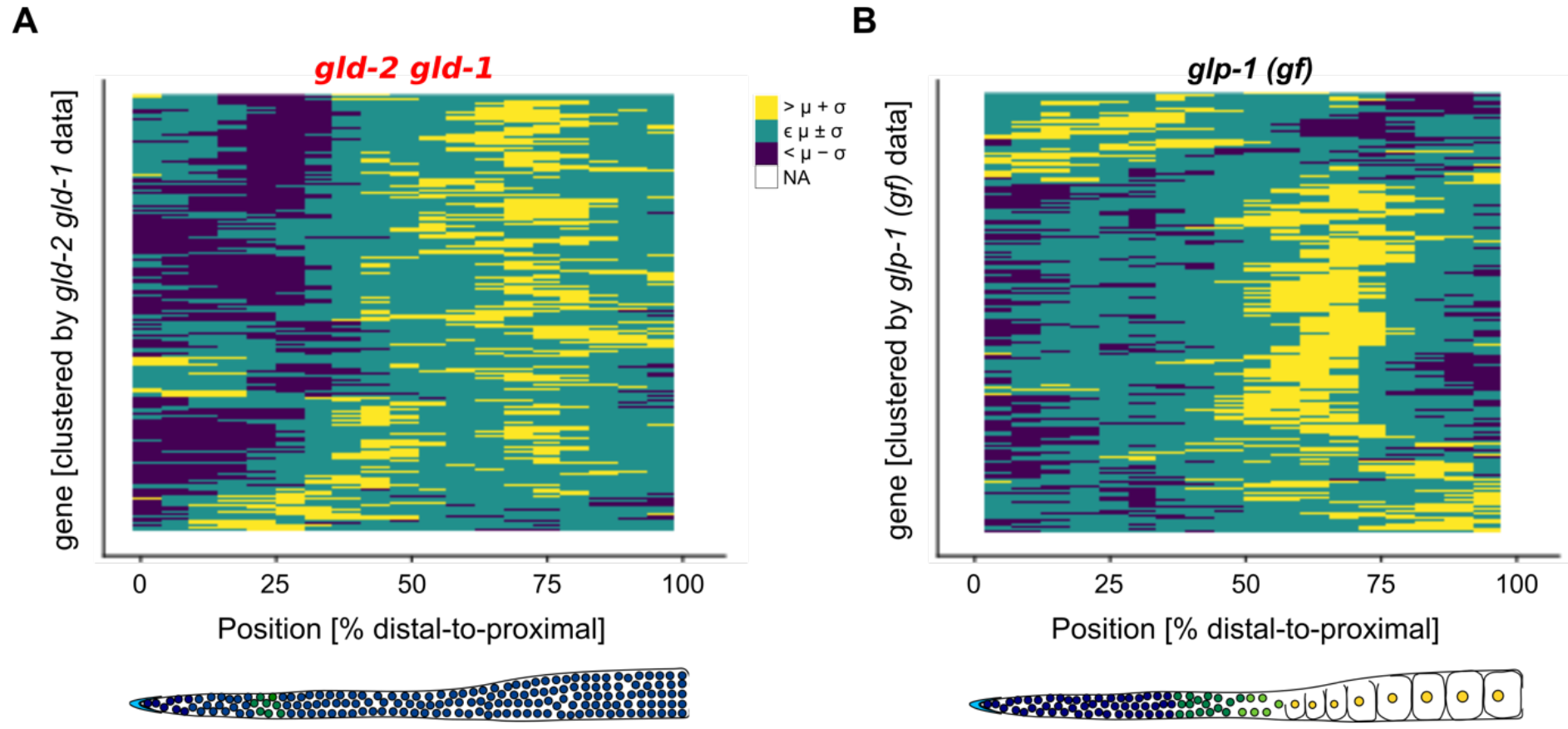
*gld-2 gld-1* double mutant and *glp-1 (gf)* mutant display mRNA localization. Related to Figure 3. (A) Hierarchical clustering of germline specific genes by linear correlation (1 - Pearson’s, r) for *gld-2 gld-1* double mutant. μ: Mean. σ: Standard deviation. NA: No data. (B) Hierarchical clustering of germline specific genes by linear correlation (1 - Pearson’s, r) for *glp-1 (gf)* mutant. μ: Mean. σ: Standard deviation. NA: no data.

**Figure S4.**
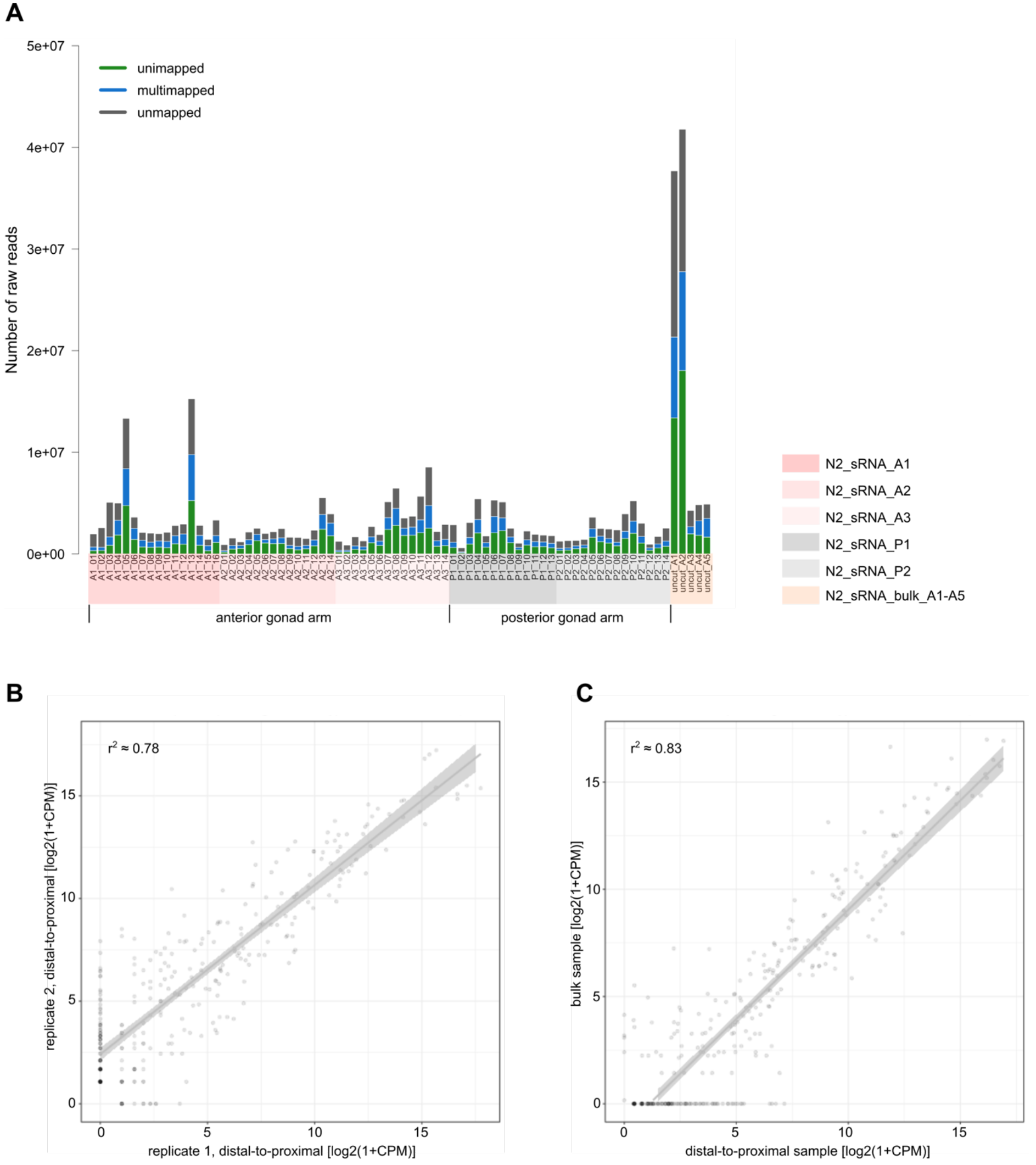
Experimental approach for spatial miRNA expression is highly reproducible and reliable. Related to Figure 1 and 4. (A) Read counts and assignability of reads for each biological and technical replicate. (B) Linear correlation (Pearson’s r) across all miRNAs of *in silico* pooled slices for two biological replicates. (C) Linear correlation (Pearson’s r) across all miRNAs of uncut (bulk) sample and *in silico* pooled slices.

**Figure S5.**
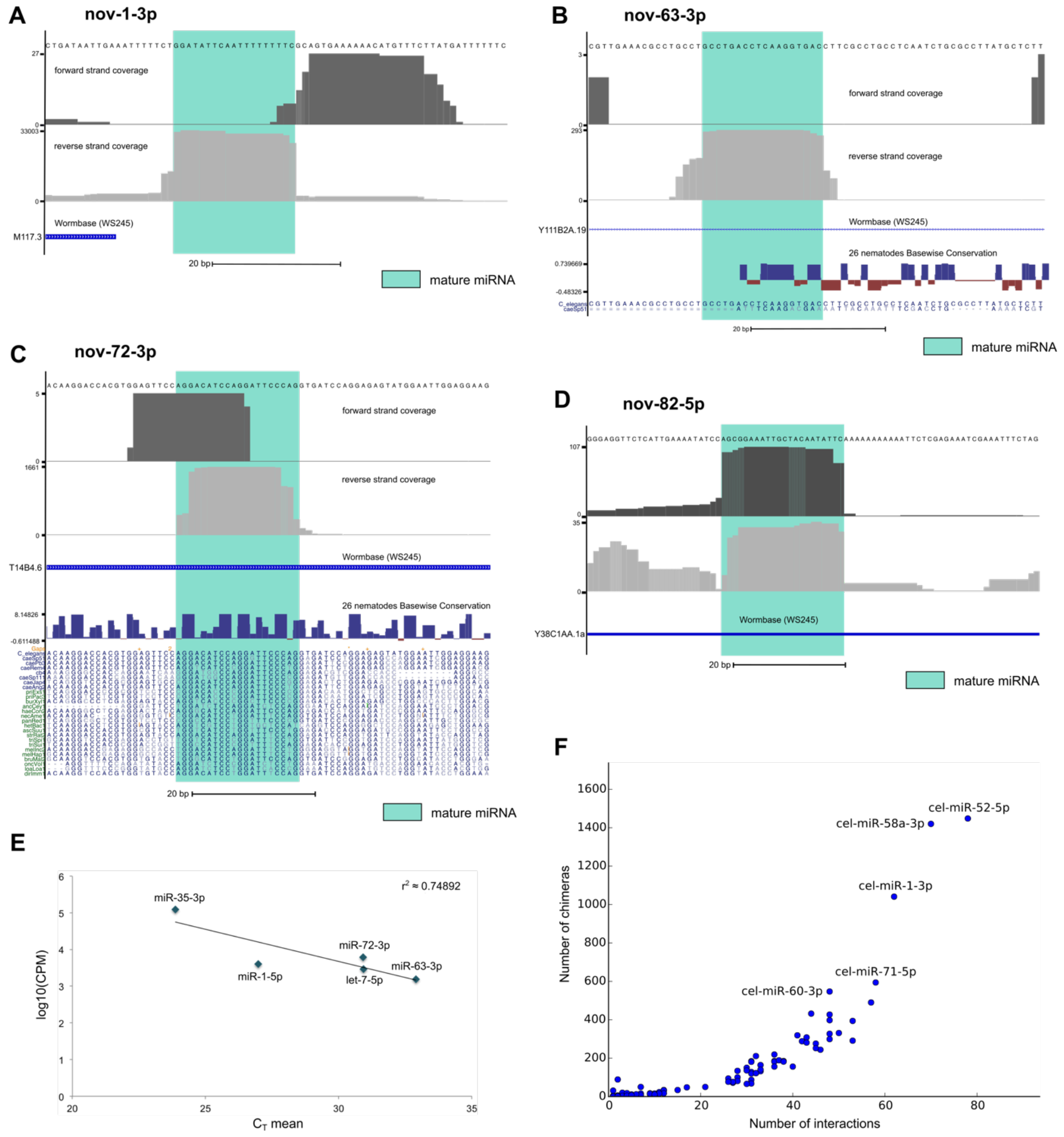
Novel miRNA predictions exhibit miRNA-like features. Related to Figure 4. (A) Genome browser track showing read coverage of predicted miRNA candidate, nov-1-3p with reads stacking up mostly on the mature sequence (light blue) at aligned 5’ positions. Forward strand coverage is indicated in dark grey and reverse strand coverage is indicated in light grey. (B) Genome browser track showing read coverage of predicted novel miRNA candidate, nov-64 with reads stacking up mostly on the mature sequence (light blue) at aligned 5’ positions. Forward strand coverage is indicated in dark grey and reverse strand coverage is indicated in light grey. Conservation across different species is displayed at nucleotide resolution. (C) Genome browser track showing read coverage of predicted novel miRNA candidate, nov-72-3p with reads stacking up mostly on the mature sequence (light blue) at aligned 5’ positions. Forward strand coverage is indicated in dark grey and reverse strand coverage is indicated in light grey. Conservation across different species is displayed at nucleotide resolution. (D) Genome browser track showing read coverage of predicted novel miRNA candidate, nov-82-5p with reads stacking up mostly on the mature sequence (light blue) at aligned 5’ positions. Forward strand coverage is indicated in dark grey and reverse strand coverage is indicated in light grey. (E) Correlation of expression (CPM) of known miRNAs (mir-35-3p, mir-1-3p and let-7-5p) and novel miRNA predictions (nov-63-3p and nov-72-3p) with corresponding C_T_ values measured by TaqMan^®^ assay (expression of mature miRNAs). (F) Number of miRNA:mRNA chimeras for the novel miRNA nov-72-3p.

**Figure S6.**
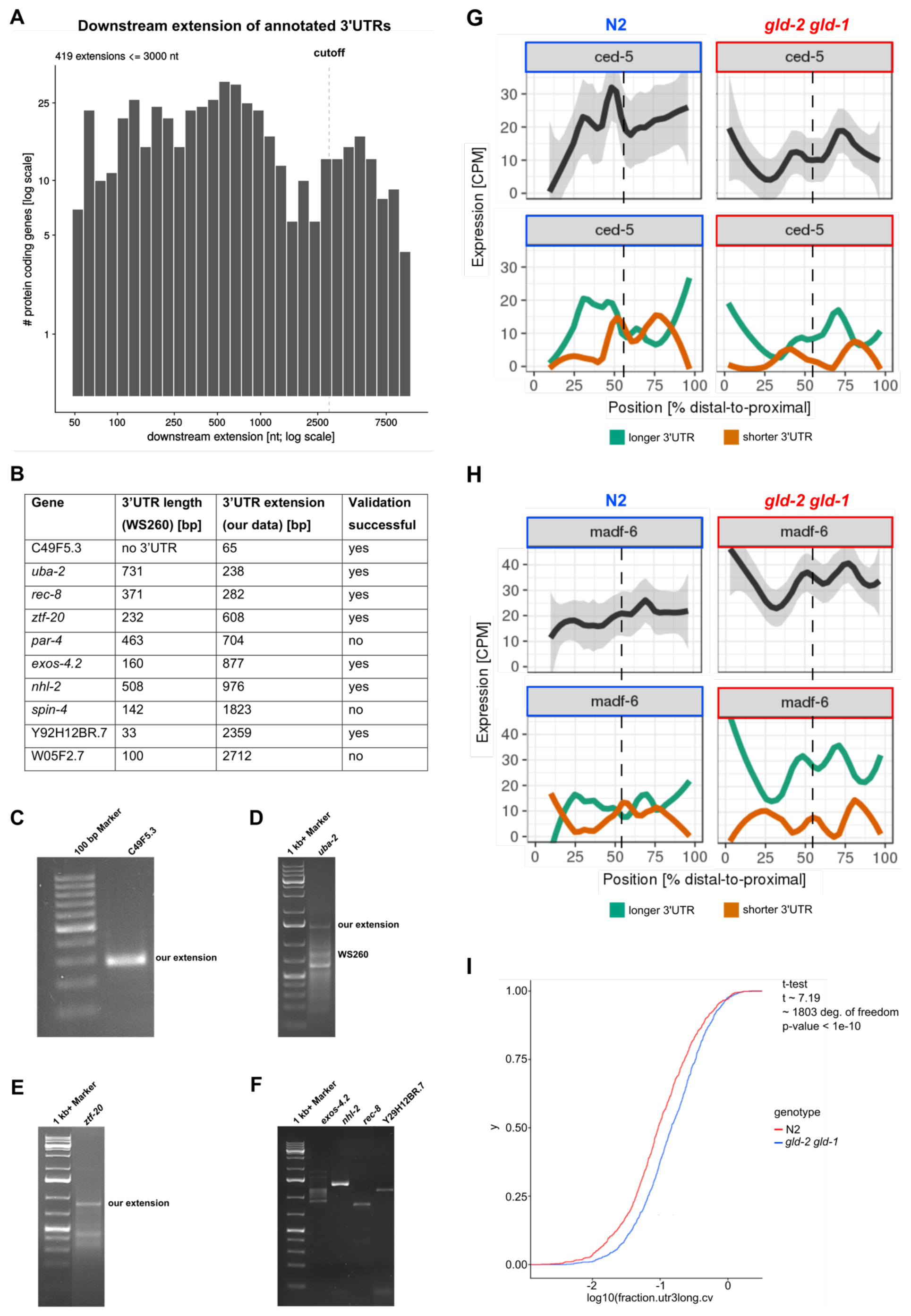
Downstream extension and validation of annotated 3’ UTRs and examples of differential 3’ UTR isoform usage across germline. Related to Figure 6. (A) Summary of all downstream extensions of annotated 3’ UTRs. Only candidates with an extension smaller or equal to 3000 nt were considered for further analysis and validation. (B) Table summarizing candidates that were chosen for downstream extension validation with annotated 3’ UTR length (WS260), downstream extension and result of validation. (C) Validation of downstream extension of C49F5.3 annotated 3’ UTR by nested PCR. Marker: 1 kb+ gene ruler. (D) Validation of downstream extension of *uba-2* annotated 3’ UTR by nested PCR. Marker: 1 kb+ gene ruler. Our extension and WS260 annotation are indicated. (E) Validation of downstream extension of *ztf-20* annotated 3’ UTR by nested PCR. Marker: 1 kb+ gene ruler. (F) Validation of downstream extension of *exos-4.2, nhl-2, rec-8* and Y29H12BR.7 annotated 3’ UTR by conventional PCR. Marker: 1 kb+ gene ruler. (G) Spatial expression of *ced-5* in wild type N2 and *gld-2 gld-1* double mutant from distal-to-proximal at gene and isoform level. n=6 independent experiments for N2 and n=4 for *gld-2 gld-1*, LOESS ± standard error (SE) for gene level and LOESS only for isoform level. Longest 3’ UTR is marked in turquoise and shorter 3’ UTR in orange. Dashed line marks the bend/loop region of the germline. (H) Spatial expression of *madf-6* in wild type N2 and *gld-2 gld-1* double mutant from distal-to-proximal at gene and isoform level. n=6 independent experiments for N2 and n=4 for *gld-2 gld-1*, LOESS ± SE for gene level and LOESS only for isoform level. Longest 3’ UTR is marked in turquoise and shorter 3’ UTR in orange. Dashed line marks the bend/loop region of the germline. (I) Comparison of the cumulative densities of 3’ UTR variability distribution between N2 and *gld-2 gld-1* double mutant. 3’ UTR variability was measured by the coefficients of variation (CV) of the contribution of longer 3’ UTR to the total expression of the top two expressed (on average) isoforms per gene. 919 genes with several isoforms expressed at 5 CPM or higher on average in either condition were considered for the analysis. 9 genes with CV’s below the 0.1^st^ percentile of the log normal fit in either condition were excluded from the analysis.

**Figure S7.**
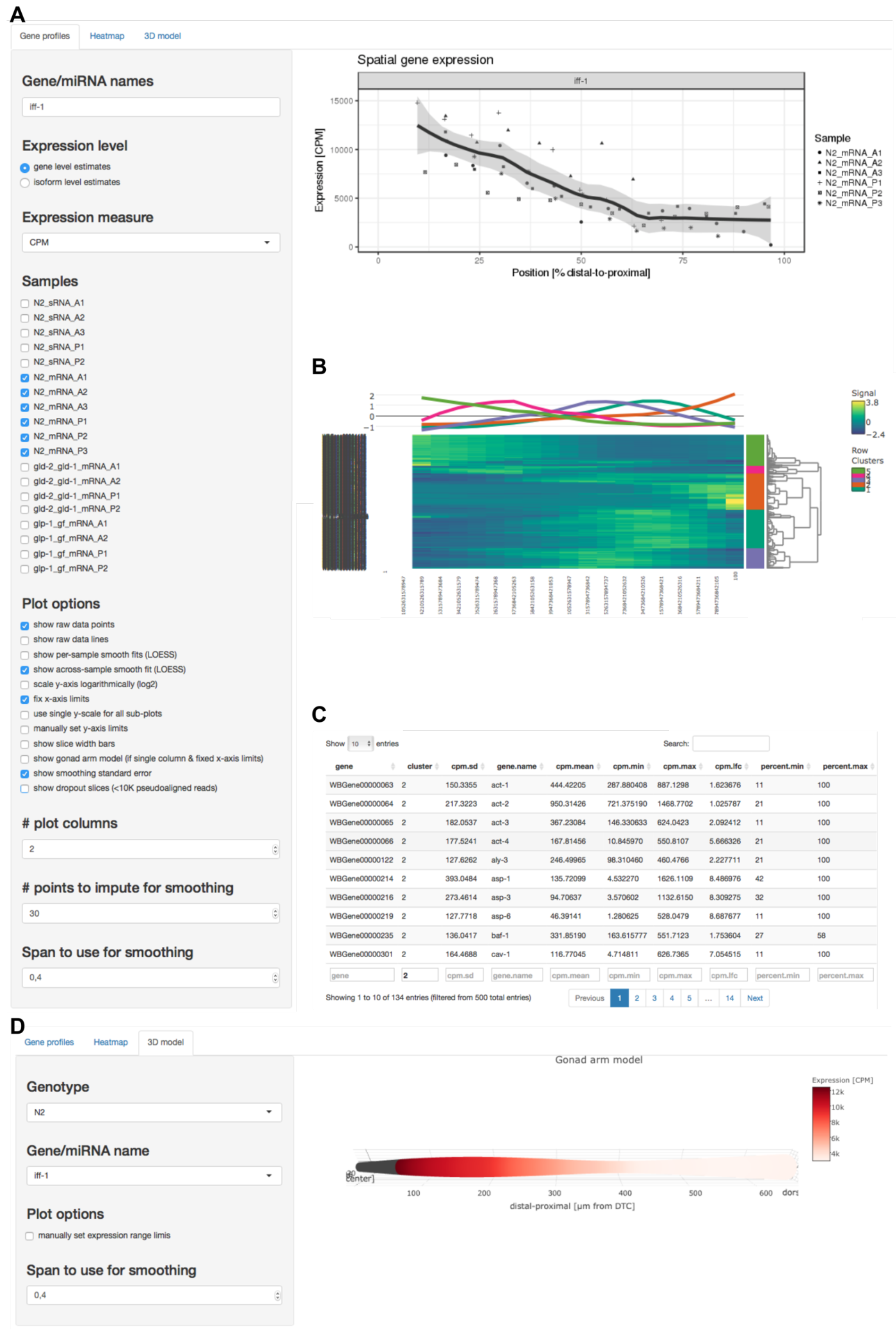
SPACEGERM: a user-friendly interface for exploring spatial expression across the germline in 2D and 3D. Related to Figure 1, 2, 3, 4 and 6. (A) Plotting options for each transcript detected in our data. As an example, the spatial expression of *iff-1* is shown for all biological and technical replicates of N2, LOESS ± standard error (SE). (B) Global spatial gene expression can be investigated by clustering all detected genes according linear correlation (Pearson, r) for all genotypes. μ: Mean. σ: standard deviation. NA: No data. (C) Result of clustering can be exported as an Excel file and investigated in more detail. (D) Virtual *in situ* hybridization (vISH) using reconstructed 3D germline model. As an example, the spatial expression of *iff-1* is shown for all biological and technical replicates of N2.

## REFERENCES

Bartel, D.P. (2009). MicroRNAs: target recognition and regulatory functions. Cell 136, 215–233.

Bartel, D.P. (2018). Metazoan MicroRNAs. Cell 173, 20–51.

Batchelder, C., Dunn, M.A., Choy, B., Suh, Y., Cassie, C., Shim, E.Y., Shin, T.H., Mello, C., Seydoux, G., and Blackwell, T.K. (1999). Transcriptional repression by the Caenorhabditis elegans germ-line protein PIE-1. Genes Dev. 13, 202–212.

Brenner, J.L., and Schedl, T. (2016). Germline Stem Cell Differentiation Entails Regional Control of Cell Fate Regulator GLD-1 in Caenorhabditis elegans. Genetics 202, 1085–1103.

Brumbaugh, J., Di Stefano, B., Wang, X., Borkent, M., Forouzmand, E., Clowers, K.J., Ji, F., Schwarz, B.A., Kalocsay, M., Elledge, S.J., et al. (2018). Nudt21 controls cell fate by connecting alternative polyadenylation to chromatin signaling. Cell 172, 106–120.e21.

Bukhari, S.I.A., Vasquez-Rifo, A., Gagné, D., Paquet, E.R., Zetka, M., Robert, C., Masson, J.-Y., and Simard, M.J. (2012). The microRNA pathway controls germ cell proliferation and differentiation in C. elegans. Cell Res. 22, 1034–1045.

Buxbaum, A.R., Haimovich, G., and Singer, R.H. (2015). In the right place at the right time: visualizing and understanding mRNA localization. Nat. Rev. Mol. Cell Biol. 16, 95–109.

Bystrykh, L.V. (2012). Generalized DNA barcode design based on Hamming codes. PLoS One 7, e36852.

Crittenden, S.L., Leonhard, K.A., Byrd, D.T., and Kimble, J. (2006). Cellular analyses of the mitotic region in the Caenorhabditis elegans adult germ line. Mol. Biol. Cell 17, 3051–3061.

D’Agostino, I., Merritt, C., Chen, P.-L., Seydoux, G., and Subramaniam, K. (2006). Translational repression restricts expression of the C. elegans Nanos homolog NOS-2 to the embryonic germline. Dev. Biol. 292, 244–252.

Dard-Dascot, C., Naquin, D., d’Aubenton-Carafa, Y., Alix, K., Thermes, C., and van Dijk, E. (2018). Systematic comparison of small RNA library preparation protocols for next-generation sequencing. BMC Genomics 19, 118.

Ding, X.C., Slack, F.J., and Grosshans, H. (2008). The let-7 microRNA interfaces extensively with the translation machinery to regulate cell differentiation. Cell Cycle 7, 3083–3090.

Evsikov, A.V., Graber, J.H., Brockman, J.M., Hampl, A., Holbrook, A.E., Singh, P., Eppig, J.J., Solter, D., and Knowles, B.B. (2006). Cracking the egg: molecular dynamics and evolutionary aspects of the transition from the fully grown oocyte to embryo. Genes Dev. 20, 2713–2727.

Fox, P.M., Vought, V.E., Hanazawa, M., Lee, M.-H., Maine, E.M., and Schedl, T. (2011). Cyclin E and CDK-2 regulate proliferative cell fate and cell cycle progression in the C. elegans germline. Development 138, 2223–2234.

Francis, R., Maine, E., and Schedl, T. Analysis of the Multiple Roles of gld-I in Germline Development: Interactions.

Friedländer, M.R., Mackowiak, S.D., Li, N., Chen, W., and Rajewsky, N. (2012). miRDeep2 accurately identifies known and hundreds of novel microRNA genes in seven animal clades. Nucleic Acids Res. 40, 37–52.

Gartner, A., Boag, P.R., and Blackwell, T.K. (2008). Germline survival and apoptosis. WormBook 1–20.

Grosswendt, S., Filipchyk, A., Manzano, M., Klironomos, F., Schilling, M., Herzog, M., Gottwein, E., and Rajewsky, N. (2014). Unambiguous identification of miRNA:target site interactions by different types of ligation reactions. Mol. Cell 54, 1042–1054.

Hansen, D., and Schedl, T. (2013). Stem cell proliferation versus meiotic fate decision in Caenorhabditis elegans. Adv. Exp. Med. Biol. 757, 71–99.

Hashimshony, T., Wagner, F., Sher, N., and Yanai, I. (2012). CEL-Seq: single-cell RNA-Seq by multiplexed linear amplification. Cell Rep. 2, 666–673.

Hashimshony, T., Senderovich, N., Avital, G., Klochendler, A., de Leeuw, Y., Anavy, L., Gennert, D., Li, S., Livak, K.J., Rozenblatt-Rosen, O., et al. (2016). CEL-Seq2: sensitive highly-multiplexed single-cell RNA-Seq. Genome Biol. 17, 77.

Hirsh, D., Oppenheim, D., and Klass, M. (1976). Development of the reproductive system of Caenorhabditis elegans. Dev. Biol. 49, 200–219.

Hubbard, E.J.A. (2007). Caenorhabditis elegans germ line: a model for stem cell biology. Dev. Dyn. 236, 3343–3357.

Jan, C.H., Friedman, R.C., Ruby, J.G., and Bartel, D.P. (2011). Formation, regulation and evolution of Caenorhabditis elegans 3’UTRs. Nature 469, 97–101.

Jansen, R.P. (2001). mRNA localization: message on the move. Nat. Rev. Mol. Cell Biol. 2, 247–256.

Jones, A.R., Francis, R., and Schedl, T. (1996). GLD-1, a cytoplasmic protein essential for oocyte differentiation, shows stage- and sex-specific expression during Caenorhabditis elegans germline development. Dev. Biol. 180, 165–183.

Jungkamp, A.-C., Stoeckius, M., Mecenas, D., Grün, D., Mastrobuoni, G., Kempa, S., and Rajewsky, N. (2011). In vivo and transcriptome-wide identification of RNA binding protein target sites. Mol. Cell 44, 828–840.

Junker, J.P., Noël, E.S., Guryev, V., Peterson, K.A., Shah, G., Huisken, J., McMahon, A.P., Berezikov, E., Bakkers, J., and van Oudenaarden, A. (2014). Genome-wide RNA Tomography in the zebrafish embryo. Cell 159, 662–675.

Kadyk, L.C., and Kimble, J. (1998). Genetic regulation of entry into meiosis in Caenorhabditis elegans. Development 125, 1803–1813.

Kaufmann, I., Martin, G., Friedlein, A., Langen, H., and Keller, W. (2004). Human Fip1 is a subunit of CPSF that binds to U-rich RNA elements and stimulates poly(A) polymerase. EMBO J. 23, 616–626.

Kim, K.W., Wilson, T.L., and Kimble, J. (2010). GLD-2/RNP-8 cytoplasmic poly(A) polymerase is a broad-spectrum regulator of the oogenesis program. Proc. Natl. Acad. Sci. USA 107, 17445–17450.

Koller, M., and Stahel, W.A. (2011). Sharpening Wald-type inference in robust regression for small samples. Comput. Stat. Data Anal. 55, 2504–2515.

Lackford, B., Yao, C., Charles, G.M., Weng, L., Zheng, X., Choi, E.-A., Xie, X., Wan, J., Xing, Y., Freudenberg, J.M., et al. (2014). Fip1 regulates mRNA alternative polyadenylation to promote stem cell self-renewal. EMBO J. 33, 878–889.

Lai, E.C., Wiel, C., and Rubin, G.M. (2004). Complementary miRNA pairs suggest a regulatory role for miRNA:miRNA duplexes. RNA 10, 171–175.

Maciejowski, J., Ugel, N., Mishra, B., Isopi, M., and Hubbard, E.J.A. (2006). Quantitative analysis of germline mitosis in adult C. elegans. Dev. Biol. 292, 142–151.

Mangone, M., Manoharan, A.P., Thierry-Mieg, D., Thierry-Mieg, J., Han, T., Mackowiak, S.D., Mis, E., Zegar, C., Gutwein, M.R., Khivansara, V., et al. (2010). The landscape of C. elegans 3’UTRs. Science (80-.). 329, 432–435.

Martin, K.C., and Ephrussi, A. (2009). mRNA localization: gene expression in the spatial dimension. Cell 136, 719–730.

Mayr, C. (2017). Regulation by 3’-Untranslated Regions. Annu. Rev. Genet. 51, 171–194.

Mayr, C., and Bartel, D.P. (2009). Widespread shortening of 3’UTRs by alternative cleavage and polyadenylation activates oncogenes in cancer cells. Cell 138, 673–684.

McEwen, T.J., Yao, Q., Yun, S., Lee, C.-Y., and Bennett, K.L. (2016). Small RNA in situ hybridization in Caenorhabditis elegans, combined with RNA-seq, identifies germline-enriched microRNAs. Dev. Biol. 418, 248–257.

Mello, C.C., Schubert, C., Draper, B., Zhang, W., Lobel, R., and Priess, J.R. (1996). The PIE-1 protein and germline specification in C. elegans embryos. Nature 382, 710–712.

Melo, J.A., and Ruvkun, G. (2012). Inactivation of conserved C. elegans genes engages pathogen- and xenobiotic-associated defenses. Cell 149, 452–466.

Merritt, C., Rasoloson, D., Ko, D., and Seydoux, G. (2008). 3’ UTRs are the primary regulators of gene expression in the C. elegans germline. Curr. Biol. 18, 1476–1482.

Millonigg, S., Minasaki, R., Nousch, M., Novak, J., and Eckmann, C.R. (2014). GLD-4-mediated translational activation regulates the size of the proliferative germ cell pool in the adult C. elegans germ line. PLoS Genet. 10, e1004647.

Mukherji, S., Ebert, M.S., Zheng, G.X.Y., Tsang, J.S., Sharp, P.A., and van Oudenaarden, A. (2011). MicroRNAs can generate thresholds in target gene expression. Nat. Genet. 43, 854–859.

Nousch, M., and Eckmann, C.R. (2013). Translational control in the Caenorhabditis elegans germ line. Adv. Exp. Med. Biol. 757, 205–247.

Nousch, M., Yeroslaviz, A., Habermann, B., and Eckmann, C.R. (2014). The cytoplasmic poly(A) polymerases GLD-2 and GLD-4 promote general gene expression via distinct mechanisms. Nucleic Acids Res. 42, 11622–11633.

Nousch, M., Minasaki, R., and Eckmann, C.R. (2017). Polyadenylation is the key aspect of GLD-2 function in C. elegans. RNA 23, 1180–1187.

Rybak-Wolf, A., Jens, M., Murakawa, Y., Herzog, M., Landthaler, M., and Rajewsky, N. (2014). A variety of dicer substrates in human and C. elegans. Cell 159, 1153–1167.

Sambrook, J., and Russell, D.W. (2006). Rapid Amplification of 3’ cDNA Ends (3’-RACE). CSH Protoc 2006.

Sandberg, R., Neilson, J.R., Sarma, A., Sharp, P.A., and Burge, C.B. (2008). Proliferating cells express mRNAs with shortened 3’ untranslated regions and fewer microRNA target sites. Science (80-.). 320, 1643–1647.

Seydoux, G., and Dunn, M.A. (1997). Transcriptionally repressed germ cells lack a subpopulation of phosphorylated RNA polymerase II in early embryos of Caenorhabditis elegans and Drosophila melanogaster. Development 124, 2191–2201.

Seydoux, G., and Fire, A. (1994). Soma-germline asymmetry in the distributions of embryonic RNAs in Caenorhabditis elegans. Development 120, 2823–2834.

Seydoux, G., Mello, C.C., Pettitt, J., Wood, W.B., Priess, J.R., and Fire, A. (1996). Repression of gene expression in the embryonic germ lineage of C. elegans. Nature 382, 713–716.

Shepard, P.J., Choi, E.-A., Lu, J., Flanagan, L.A., Hertel, K.J., and Shi, Y. (2011). Complex and dynamic landscape of RNA polyadenylation revealed by PAS-Seq. RNA 17, 761–772.

Sood, P., Krek, A., Zavolan, M., Macino, G., and Rajewsky, N. (2006). Cell-type-specific signatures of microRNAs on target mRNA expression. Proc. Natl. Acad. Sci. USA 103, 2746–2751.

Stoeckius, M., Grün, D., Kirchner, M., Ayoub, S., Torti, F., Piano, F., Herzog, M., Selbach, M., and Rajewsky, N. (2014). Global characterization of the oocyte-to-embryo transition in Caenorhabditis elegans uncovers a novel mRNA clearance mechanism. EMBO J. 33, 1751–1766.

Tenenhaus, C., Subramaniam, K., Dunn, M.A., and Seydoux, G. (2001). PIE-1 is a bifunctional protein that regulates maternal and zygotic gene expression in the embryonic germ line of Caenorhabditis elegans. Genes Dev. 15, 1031–1040.

Wang, X., Zhao, Y., Wong, K., Ehlers, P., Kohara, Y., Jones, S.J., Marra, M.A., Holt, R.A., Moerman, D.G., and Hansen, D. (2009). Identification of genes expressed in the hermaphrodite germ line of C. elegans using SAGE. BMC Genomics 10, 213.

West, S.M., Mecenas, D., Gutwein, M., Aristizábal-Corrales, D., Piano, F., and Gunsalus, K.C. (2018). Developmental dynamics of gene expression and alternative polyadenylation in the Caenorhabditis elegans germline. Genome Biol. 19, 8.

Wolke, U., Jezuit, E.A., and Priess, J.R. (2007). Actin-dependent cytoplasmic streaming in C. elegans oogenesis. Development 134, 2227–2236.

(2009). Correction for Ji et al., Progressive lengthening of 3’ untranslated regions of mRNAs by alternative polyadenylation during mouse embryonic development. Proc. Natl. Acad. Sci. USA 106, 9535–9535.

